# Unraveling the Neurocognitive Mechanisms of Delayed Punishment in Second-and Third-Party Contexts

**DOI:** 10.1101/2025.02.25.640024

**Authors:** Jiamin Huang, Yuwen He, Zejian Chen, Qianwei Cai, Yantong Yao, Yuanchen Wang, Yanyan Qi, Haiyan Wu

## Abstract

Second-party punishment (SPP) and third-party punishment (TPP) are essential in regulating social behavior and maintaining social norms, yet the effectiveness may wane when punishment is delayed. Currently, the decision-making and neural mechanisms underlying SPP and TPP under temporal delays remain largely unexplored. This study investigated both SPP and TPP punishment decisions with time-delay (immediate vs. delayed) using a hypothetical criminal scenario assessment task with fMRI. Results showed increased activity in the bilateral precuneus and left temporoparietal junction (TPJ) and stronger left TPJ-dorsolateral prefrontal cortex (dlPFC) connectivity in SPP compared to TPP. Interestingly, third-parties imposed more severe punishment in delayed conditions than in immediate ones, accompanied by enhanced neural activity in dorsomedial prefrontal cortex (dmPFC), left TPJ, ventrolateral prefrontal cortex (vlPFC), ventromedial prefrontal cortex (vmPFC), and caudate nucleus. This delay effect was evident only in misdemeanor and felony cases, with no differences in capital offenses, suggesting a moderating effect of criminal severity. In contrast, SPP showed no significant changes in punishment or neural response across immediate and delayed conditions. Multivariate pattern analysis further indicated that dmPFC, TPJ, vlPFC, vmPFC and caudate function together to encode punishment severity in TPP contexts, a pattern not observed in SPP. These findings underscore distinct decision-making mechanisms between SPP and TPP under temporal delays, with implications for understanding justice-related processing in the human brain under time constraints.

**Significance statement:** Within the justice system, punishing norm violators is often hindered when they flee, resulting in delayed punishments. However, little is known about how second-party (SPP) and third-party (TPP) implement punishment with time delays. The current study investigated this using real criminal scenarios with time delays under fMRI scanning. We found that third parties imposed more severe punishment on defendants in delayed conditions than in immediate ones, reflected by enhanced neural activity in the dorsomedial prefrontal cortex, left temporoparietal junction, ventrolateral prefrontal cortex, ventromedial prefrontal cortex, and caudate nucleus. In contrast, no significant differences were observed in second-party punishment. We also found a moderating effect of criminal severity: the difference between immediate and delayed sanctions by third-parties was present in misdemeanor and felony cases but not in capital offenses. These results shed light on the decision-making mechanisms of SPP and TPP in contexts involving temporal delay, which may have deep impact in different disciplines, e.g., social science, cognitive neuroscience and legal system.

## Introduction

In human society, both second-party punishment (SPP) and third-party punishment (TPP) are integral to the enforcement of social norms (Fehr & Fischbacher, 2004; Henrich & Muthukrishna, 2021). SPP is often driven by a desire for retribution or the expression of personal feelings, enacted by individuals directly impacted by norm violations. Conversely, TPP is carried out by observers, focusing not only on the harm inflicted on the victim but also on the intentions of the wrongdoer and the broader implications for social norms or laws (Buckholtz & Marois, 2012; Fehr & Fischbacher, 2004; Mohlin et al., 2023). Several studies comparing the differences between SPP and TPP have indicated TPP might be more impartial and be able to consider the contextual information for more comprehensive decision-making (Buckholtz et al., 2008; Buckholtz & Marois, 2012; Fehr & Fischbacher, 2004; Feng et al., 2022; Stallen et al., 2018). However, existing studies on SPP or TPP considered little of the timing of punishment. Especially, in real cases, punishment implementation was often hindered as the perpetrator flee, which greatly reduces the efficiency of execution. Thus, research is needed to explore the decision-making process that considers the punishment timing within TPP and SPP contexts.

Substantial work has advanced our understanding of the neural mechanisms underlying punishment decision-makings. According to the hierarchical punishment model (HPM) proposed by researchers (Krueger & Hoffman, 2016; Yang et al., 2024) as well as recent metal-analytic validation of the model (Bellucci et al., 2020), a hierarchical brain network is involved in punishment decision-making. Firstly, the salience network (SN) detects social norm violations and generates an aversive experience through anterior cingulate cortex (ACC) and anterior insula (AI), signaling harm severity via the amygdala. Subsequently, the default mode network (DMN) integrates harm and intent through self-referential and other-inferring processes, leading to an assessment of blame in ventromedial prefrontal cortex (vmPFC), dorsomedial prefrontal cortex (dmPFC), posterior cingulate cortex (PCC), and temporoparietal junction (TPJ). Finally, the central executive network (CEN), including the bilateral dorsolateral prefrontal cortex (dlPFC) and posterior parietal cortex (PPC), guides the imposition of punishment. Additionally, the reward system is implicated, as upholding social fairness by punishing norm violators is often perceived as rewarding (de Quervain et al., 2004; Stallen et al., 2018; Strobel et al., 2011). Although this pathway has been well-documented, the neural processes underlying delayed punishment remain rarely understood, particularly regarding how activations may differ between SPP and TPP.

The current study aimed to explore the behaviors and neural circuits underlying immediate and delayed sanctions using the hypothetical criminal scenario paradigm (Fig. 1), with real cases sourced from China Judgment Document Network (https://wenshu.court.gov.cn/). This approach enhances the ecological validity of the study. Participants were randomly assigned to either the SPP group or the TPP group. In the SPP group, participants assumed the role of the victim in a legal case, while those in the TPP group took on the role of the judge. The main task was completed in the fMRI scanner.

**Fig. 1.**
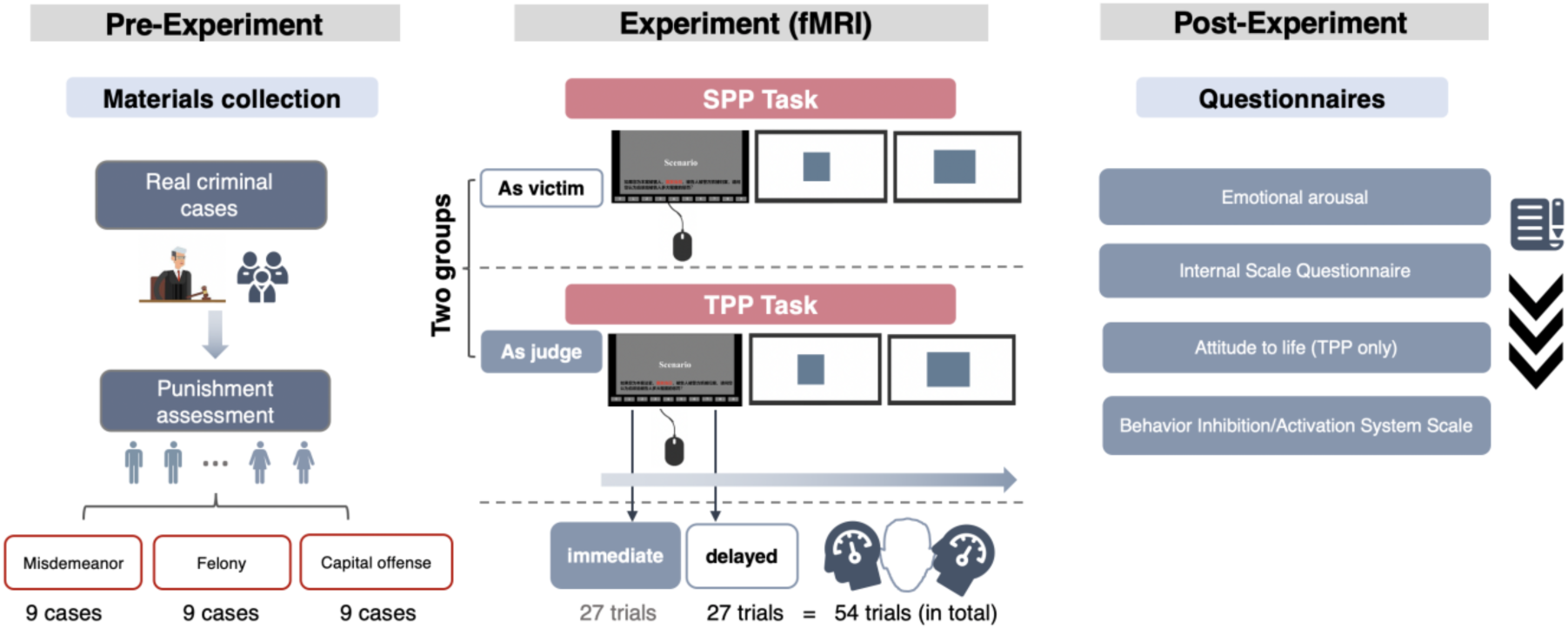
**The scheme of the experimental design**. In the fMRI task, each trial began with an infinite-time scenario displayed on the screen. This scenario remained until the participant pressed the mouse to determine the degree of punishment under the SPP (as a second party) and TPP (as a third party) tasks. Once the scenario screen disappeared, a small white square appeared for 6000 - 8000 ms. Subsequently, a large white square was shown for 2000 ms before disappearing, indicating the end of the trial and the beginning of the next one. After the fMRI task, we collected ratings from the participants.

Given that assuming the role of the victim (i.e., SPP) should evoke more personal involvement than assuming the role of the judge (i.e., TPP), we hypothesized that SPP may engage more theory of mind (ToM). When comparing decision-making differences between immediate and delayed sanctions, it is important to consider the established presence of delay discounting in value computation (Pinger et al., 2022; Volkow & Baler, 2015). Previous research has shown that delayed punishments are perceived as less severe by the first-party (perpetrator) in both human and animal studies (Bernasco et al., 2017; Liley et al., 2019). Based on these findings, we hypothesized that individuals may account for the delay discounting, leading to the imposition of harsher punishments when sanctions are delayed. The level of emotional arousal might also vary under different temporal circumstances. At the neural level, the difference between immediate and delayed sanctions is expected to be reflected primarily in the DMN. Within this network, brain regions such as the dmPFC, vmPFC, and TPJ are hypothesized to integrate the specifics of the case and the timing of the delay through self-referential and other-inferring processes to lead to a punishment decision-making. Additionally, the role of the individual’s identity (i.e., as a second or third party) and the severity of the case are expected to serve as moderating factors.

## Results

### Behavioral results

Participants in the SPP and TPP groups made punishment decisions for transgressors in criminal scenarios of varying severity and time-delays while in the scanner. After that, all participants finished the questionnaires regarding emotional arousal and life attitude after reading each criminal scenario (see Fig.1 and *Methods*). Therefore, we presented behavioral findings as below.

#### Punishment ratings

A 2 (Time delay: immediate vs. delayed) × 3 (Criminal scenario: misdemeanor vs. felony vs. capital offense) × 2 (Group: SPP vs. TPP) mixed-design ANOVA was conducted on the degree of punishment, with Time delay and Criminal scenario as within-subjects variables and Group as a between-subjects factor (Fig. 2A). The analysis revealed a significant main effect of Time delay, *F* (1, 54) = 4.568, *p* =.037, *η*_p_² =.078, with higher degrees of punishment in the delayed condition (*M* = 5.564, *SE* =.087) compared to the immediate condition (*M* = 5.415, *SE* =.082). This result suggests that participants compensated for the discounting of punishment over time. A significant main effect of Criminal scenario was also found, *F* (2, 108) = 1311.631, *p* <.001, *η*_p_² =.960, with post-hoc analyses showing that misdemeanor cases (*M* = 2.465, *SE* =.118) received significantly less punishment than felony cases (*M* = 5.474, *SE* =.108), and felony cases received less punishment than capital offenses (*M* = 8.529, *SE* =.079). Significant interactions between Group and Time delay, *F* (1, 54) = 10.523, *p* =.002, *η*_p_² =.163, and between Criminal scenario and Time delay, *F* (2, 108) = 3.471, *p* =.035, *η*_p_² =.060, were observed. However, the other effects were non-significant, *p* ≥.186.

**Fig. 2.**
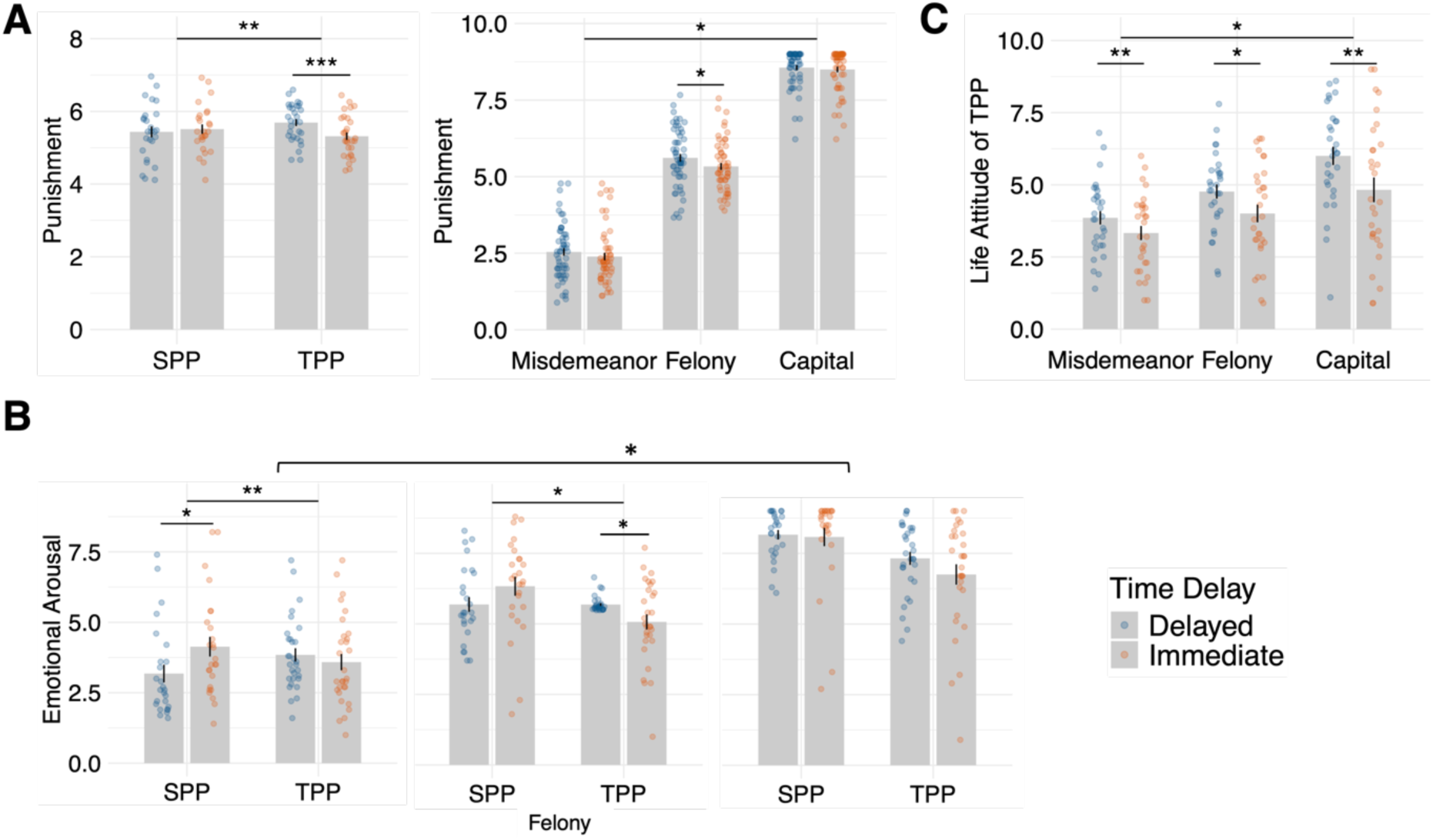
**Behavioral Results**. (A) left panel: A significant interaction of Group × Time delay on punishment ratings; right panel: A significant interaction of Criminal scenario × Time delay on punishment ratings; (B) A significant interaction of Group × Time delay × Criminal scenario on emotional arousal; (C) A significant interaction of Criminal scenario × Time delay on life attitude in the TPP group, where higher values indicate a more negative attitude. * *p* <.05, ** *p* <.01, *** *p* <.001.

Further analysis of the Group × Time delay interaction (Fig. 2A the left panel) showed that the TPP group showed significantly higher punishment in delayed condition (*M* = 5.691, *SE* =.118) compared to immediate condition (*M* = 5.317, *SE* =.112), *t* (29) = - 3.662, *p* <.001, Cohen’s *d* =.669. No significant difference was found for the SPP group, *t* (25) = 0.835, *p* =.409, Cohen’s *d* = 0.165. These findings indicate that the compensation for the discounting of the punishment only occurred in third-party context.

Regarding the Criminal scenario × Time delay interaction (Fig. 2A the right panel), participants imposed significantly higher punishment in delayed condition (*M* = 5.615, *SE* =.126) compared to immediate condition (*M* = 5.333, *SE* =.112) for felony cases, *t* (55) =-2.647, *p* =.011, Cohen’s *d* =-.796. For misdemeanors (*t* (55) =-1.919, *p* =.06, Cohen’s *d* =-.588) and capital offenses (*t* (55) =-0.803, *p* =.425, Cohen’s *d* =-.573), the differences were not significant. These findings suggested a modulating effect of Criminal scenario on punishment.

#### Post-scan emotional arousal ratings

Then we conducted the same ANOVA on emotional arousal. The results revealed a significant main effect of Group, *F* (1, 54) = 4.601, *p =*.036, *η*_p_^2^ =.079, showing a higher degree of emotional arousal in SPP group (*M* = 5.930, *SE* =.189) than that in TPP group (*M* = 5.377, *SE* =.176); a significant main effect of Criminal scenario, *F* (2, 108) = 264.951, *p <*.001, *η*_p_^2^ =.831, post hoc analysis evidenced that the degree of emotional arousal for misdemeanor (*M* = 3.686, *SE* =.184) was significantly lower than for felony (*M* = 5.695, *SE* =.125), *t* (55) =-13.857, *p* <.001, Cohen’s *d* =-1.852; and degree of emotional arousal for felony (*M* = 5.695, *SE* =.125) was significantly lower than for capital offense (*M* = 7.580, *SE* =.170), *t* (55) =-14.296, *p* <.001, Cohen’s *d* =-1.910. Further, there yielded significant two-way interactions between Criminal scenario and Time delay, *F* (2, 108) = 10.150, *p <*.001, *η*_p_^2^ =.158, between Group and Time delay, *F* (1, 54) = 5.912, *p =*.018, *η*_p_^2^ =.099, and between Group and Criminal scenario, *F* (2, 108) = 5.763, *p =*.013, *η_p_*^2^ =.096. Additionally, the three-way interaction between Group, Time delay, and Criminal scenario was significant (Fig. 2B), *F* (2, 108) = 4.171, *p =*.021, *η*_p_^2^ =.072.

Exploring the three-way interaction between Group, Time delay, and Criminal scenario revealed significant interaction between Group and Time delay in the misdemeanor scenarios (Fig. 2B), *F* (1, 54) = 9.214, *p =*.004, *η*_p_^2^ =.146. The SPP group exhibited a significantly higher level of emotional arousal in immediate condition (*M* = 4.135, *SE* =.328) compared to delayed condition (*M* = 3.181, *SE* =.283), *t* (25) = 2.505, *p* =.019, Cohen’s *d* =.491. Conversely, no significant difference was found in TPP group, *t* (29) =-1.503, *p* =.144, Cohen’s *d* =-.274. Significant interaction was also identified in the felony scenarios (Fig. 2B), *F* (1, 54) = 6.795, *p =*.012, *η*_p_^2^ =.112. Further analysis indicated that TPP group had a significantly higher degree of emotional arousal in delayed condition (*M* = 5.686, *SE* =.169) compared to immediate condition (*M* = 5.070, *SE* =.289), *t* (30) =-2.309, *p* =.028, Cohen’s *d* =-.422. No significant difference was noted for SPP group, *t* (25) = 1.540, *p* =.136, Cohen’s *d* =.302. However, the interaction in the capital offense was not significant (Fig. 2B), *F* (1, 54) = 1.225, *p =*.268, *η*_p_^2^ =.023.

#### Post-scan life attitude ratings

A 2 (Time delay: immediate vs. delayed) × 3 (Criminal scenario: misdemeanor vs. felony vs. capital offense) two-way repeated measures ANOVA was conducted to assess participants’ attitudes toward life in TPP group (Fig. 2C), where higher values indicate a more negative attitude. The analysis revealed a significant main effect of Criminal scenario, *F* (2, 58) = 29.046, *p* <.001, *η*_p_² =.50. Post-hoc analyses showed that participants’ attitudes were most negative in the capital offense (*M* = 5.415, *SE* =.333), followed by the felony cases (*M* = 4.387, *SE* =.232), *t* = 5.569, *p* <.001, Cohen’ *d* = 1.107, and were least negative in the misdemeanor (*M* = 3.593, *SE* =.221), felony vs. misdemeanor, *t* = 4.582, *p* <.001, Cohen’s *d* =.837. Additionally, there was a significant main effect of Time delay, *F* (1, 29) = 9.432, *p* =.005, *η*_p_² =.245, with attitudes being more negative in the delayed condition (*M* = 4.876, *SE* =.222) compared to the immediate condition (*M* = 4.054, *SE* =.301).

Importantly, a significant interaction between Criminal scenario and Time delay was found (Fig. 2C), *F* (2, 58) = 6.044, *p* =.014, *η*_p_² =.172, indicating that the effect of Time delay on attitudes varied depending on the criminal scenario. Simple effect analysis revealed that in misdemeanor condition, attitudes were significantly less negative in delayed condition (*M* = 3.857, *SE* =.232) compared to immediate condition (*M* = 3.330, *SE* =.245), *t* (29) =-2.906, *p* =.007, Cohen’s *d* =-.531. For felony condition, the delayed condition (*M* = 4.767, *SE* =.246) also showed less negative attitudes than the immediate condition (*M* = 4.007, *SE* =.305), *t* (29) = - 2.513, *p* =.018, Cohen’s *d* =-.459, though the effect size was smaller. The most pronounced difference was observed in capital offense condition, where attitudes were significantly less negative in the delayed condition (*M* = 6.003, *SE* =.317) compared to the immediate condition (*M* = 4.827, *SE* =.429), *t* (54) =-3.315, *p* =.002, Cohen’s *d* =-.605.

### fMRI results

#### Differences in brain responses between SPP and TPP

Confirming our hypothesis that stronger brain activations related to theory of mind would be induced in SPP than TPP, the brain-wide analysis revealed greater activation in the bilateral precuneus, with a peak in left hemisphere at MNI x/y/z =-8/-46/34 (*p*_uncorr_ <.001, *p*_FWE-corr_ =.001, see Fig. 3A) during felony criminal scenarios in SPP compared to TPP (see Table S2 for details). This finding suggests that the heightened involvement of the left precuneus reflects distinct psychological process in SPP relative to TPP.

**Fig. 3.**
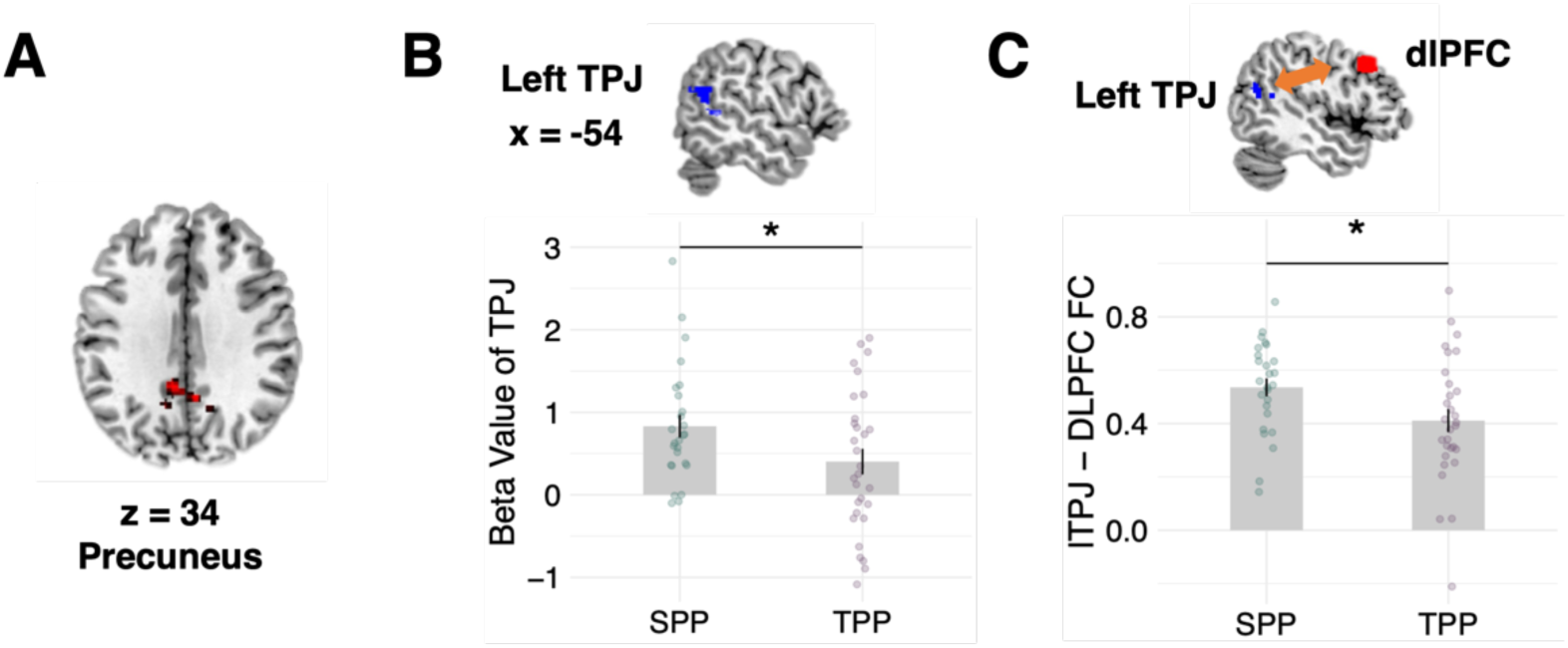
**The difference in brain responses between second-party punishment (SPP) and third-party punishment (TPP)**. (A) Activation map of the bilateral precuneus (MNI: x/y/z = - 8/-46/34) in SPP compared to TPP in the context of the felony criminal scenario in whole-brain analysis; (B) ROI analysis of left temporoparietal junction (TPJ, MNI: x/y/z =-54/-46/6) in the context of felony scenarios; (C) Task-based functional connectivity (FC) between the left TPJ (MNI: x/y/z =-54/-46/6) and the left dorsolateral prefrontal cortex (dlPFC, MNI: x/y/z =-30/38/30). * *p* <.05.

Similarly, given that the precuneus and the TPJ are both central to theory of mind, we conducted an ROI analysis focusing on the left TPJ (MNI: x/y/z =-54/-46/6). The results demonstrated that during felony scenarios, the SPP group exhibited stronger activation in the left TPJ (Fig. 3B, *t* (54) = 2.032, *p* =.047) compared to TPP group.

Integrating previous finding that showed stronger functional connectivity (FC) between the right TPJ and right dlPFC during TPP compared to SPP (Feng et al., 2022), along with our observation of stronger activation in the left TPJ during SPP compared to TPP, we conducted a FC analysis between the left TPJ and left dlPFC to assess connectivity differences between SPP and TPP in the present study. As shown in Fig. 3C, the task-based FC between the left TPJ and left dlPFC was significantly stronger in SPP than in TPP (*t* (54) = 2.242, *p* =.029).

#### Differences in brain responses between immediate and delayed sanctions

When comparing immediate and delayed conditions, whole-brain analysis (see Table S3 for details) revealed that the TPP group exhibited stronger activity in the left dmPFC (a peak at MNI: x/y/z =-14/60/26, *p*_uncorr_ = 0.001, *p*_FWE-CORR_ = 0.018) during the delayed condition compared to the immediate condition (see Fig. 4A). In contrast, the SPP group showed no significant differences.

**Fig. 4.**
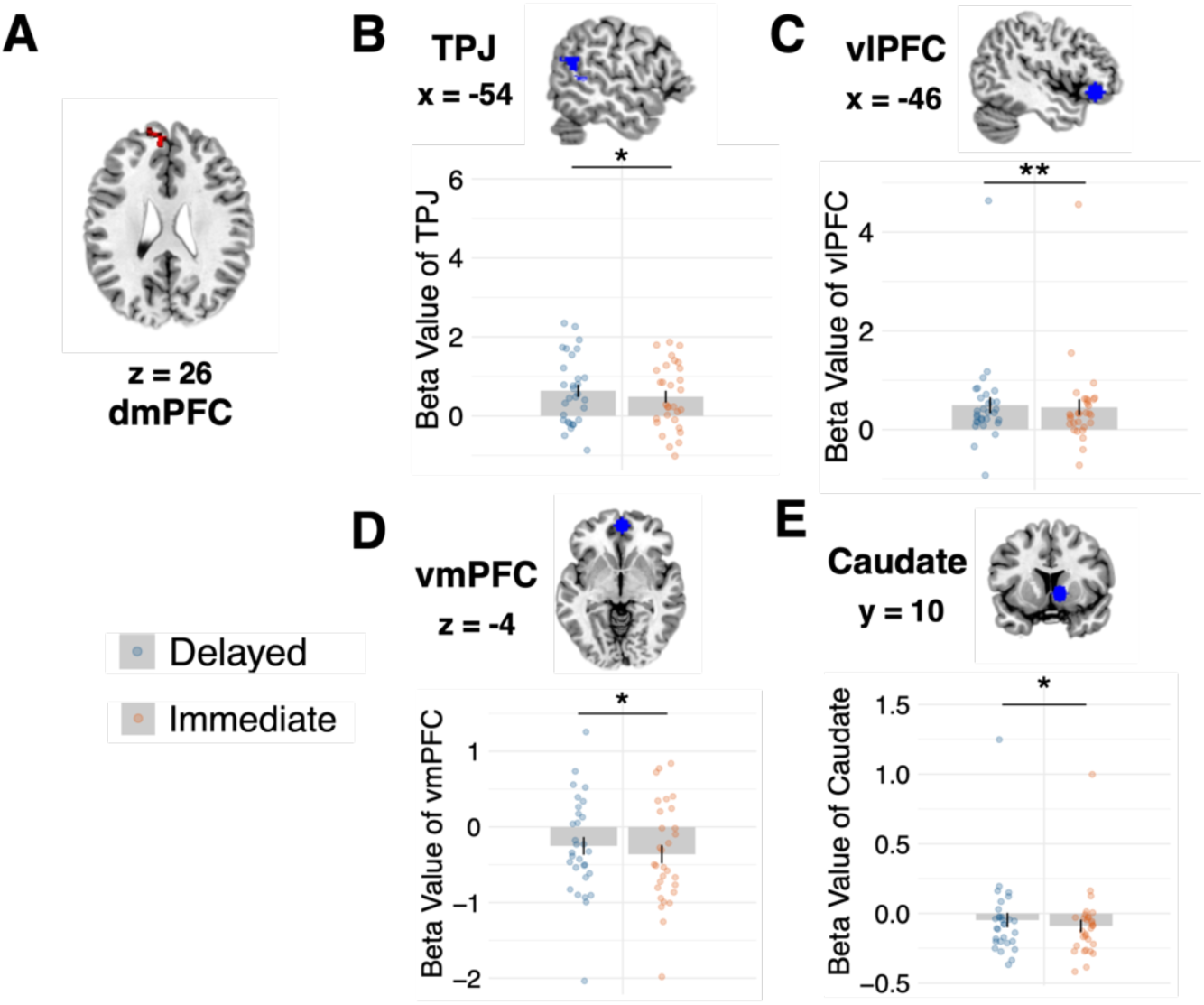
Differences in brain responses between immediate and delayed sanctions in the third-party punishment (TPP) group. (A) Activation map of the dorsomedial prefrontal cortex (dmPFC, MNI: x/y/z =-14/60/26) comparing delayed sanctions to immediate sanctions in the TPP group; (B-E) ROI analyses of the temporoparietal junction (TPJ, MNI: x/y/z =-54/-46/6), ventrolateral prefrontal cortex (vlPFC, MNI: x/y/z =-46/30/-16), ventromedial prefrontal cortex (vmPFC, MNI: x/y/z = 2/54/-4) and caudate (MNI: x/y/z = 10/10/-2) in the delayed condition compared to the immediate condition in the TPP group. * *p* <.05, ** *p* < 0.01.

In ROI analysis, we hypothesized that regions within the DMN, including dmPFC, TPJ, and vmPFC, would show stronger activation in the delayed condition compared to the immediate condition, based on the assumption that TPP requires greater integration of information than SPP. Additionally, a meta-analysis study highlighted stronger activity in the vlPFC, a key region involved in the regulation of prosocial behaviors in TPP (Bellucci et al., 2020). Furthermore, we hypothesized that brain regions associated with reward processing would exhibit enhanced activation in the delayed condition, as the implementation of punishment to uphold social fairness can be interpreted as a form of reward (de Quervain et al., 2004; Stallen et al., 2018; Strobel et al., 2011), particularly in the context of delayed sanctions. Based on these hypotheses, we selected TPJ (MNI: x/y/z =-54/-46/6), vmPFC (MNI: x/y/z = 2/54/– 4), vlPFC (MNI: x /y/z =-46/30/-16) and caudate (MNI: x/y/z = 10/10/-2) as ROIs for the analysis (see Method section for details), excluding the dmPFC due to its already identified role in the whole brain analysis. The results showed that the TPP group exhibited significantly stronger activation in the delayed condition compared to the immediate condition in TPJ (Fig. 4B), vlPFC (Fig. 4C), vmPFC (Fig. 4D), and caudate (Fig. 4E).

When analyzing cases by severity, no differences were observed between immediate and delayed conditions in capital offense cases. In felony cases (SI Appendix Fig. S2), we identified (marginally) significant activation in dmPFC (*t* (29) = 2.855, *p* =.008, *p*_FDR-corr_ =.037), TPJ (*t* (29) = 1.754, *p* =.090, *p*_FDR-corr_ =.090), vmPFC (*t*(29) = 2.307, *p* =.028, *p*_FDR-corr_ =.047), vlPFC (*t*(29) = 1.848, *p* =.0748, *p*_FDR-corr_ =.090), and Caudate (*t*(29) = 2.596, *p* =.021, *p*_FDR-corr_ =.037). In misdemeanor cases (SI Appendix Fig. S3), differences were limited to the dmPFC (*t* (29) = 2.097, *p* =.045, *p*_FDR-corr_ =.161) and TPJ (*t* (29) = 1.923, *p* =.064, *p*_FDR-corr_ =.161), though the corrected *p* values were no longer significant.

Conversely, no significant differences between immediate and delayed conditions were found in the SPP group, either across all cases or when analyzed by severity (*p* > 0.5).

#### Correlation between brain activity and punishment ratings

Given that more severe punishment was assigned in the delayed sanction, indicating punishment discounting, we conducted correlation analyses to determine whether and which brain activity could account for this phenomenon. We specifically examined the activity difference (immediate vs. delayed) in the dmPFC, TPJ, vlPFC, vmPFC, and caudate, as these regions exhibited significant activation differences across the two conditions. The correlation analysis revealed that the activation differences (*immediate – delayed*) in the TPJ (Fig. 5A, *r* =-.422, *p* =.032) and caudate (Fig. 5B, *r* =-.422, *p* =.034) were negatively correlated with the differences in punishment ratings (*immediate – delayed*) in felony criminal scenarios for the SPP group.

**Fig. 5.**
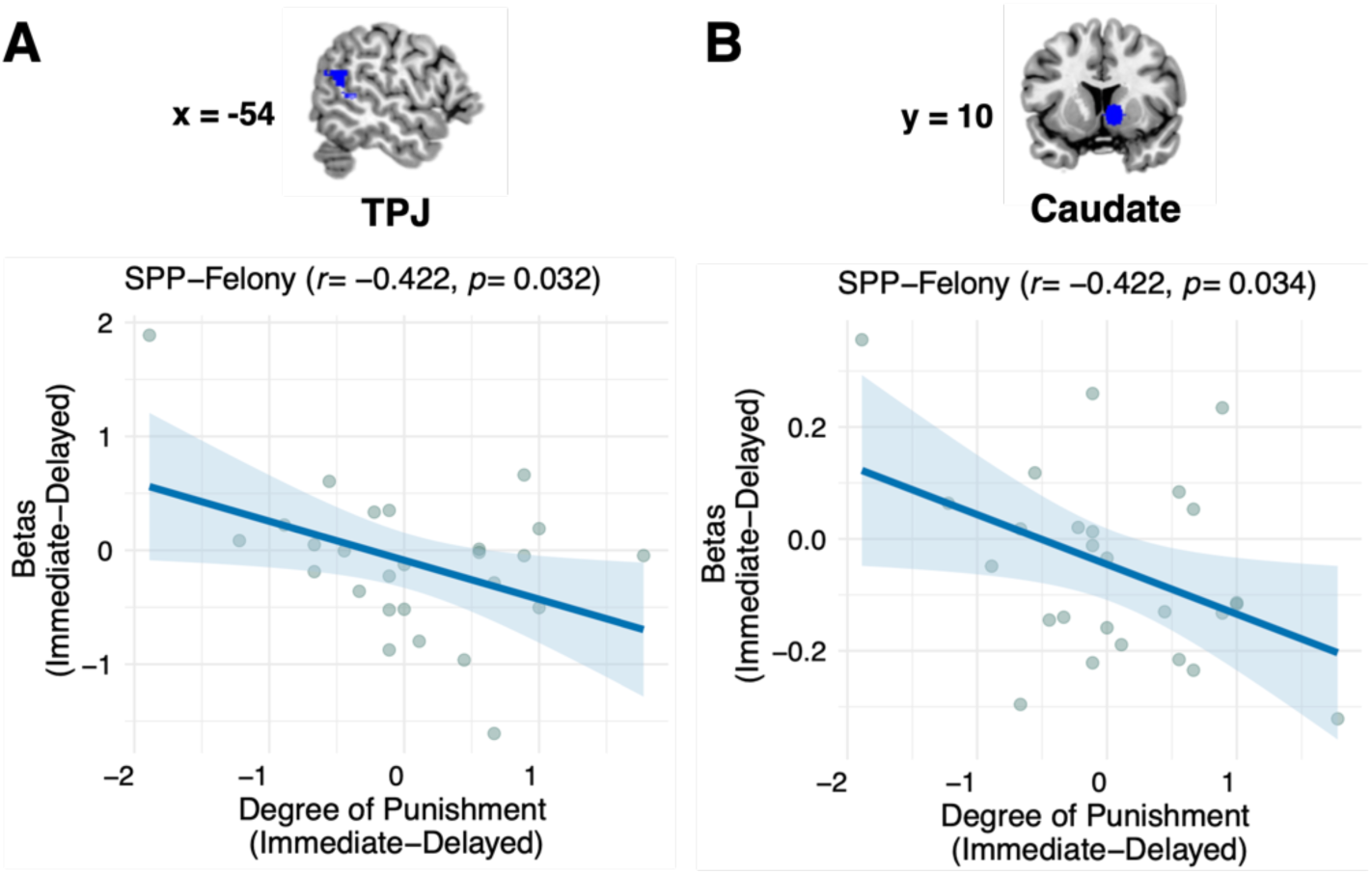
Brain activity associated with punishment ratings. The punishment differences between immediate and delayed sanctions correlates negatively with the brain activity differences between immediate and delayed conditions in (A) TPJ and (B) caudate.

**Fig. 6.**
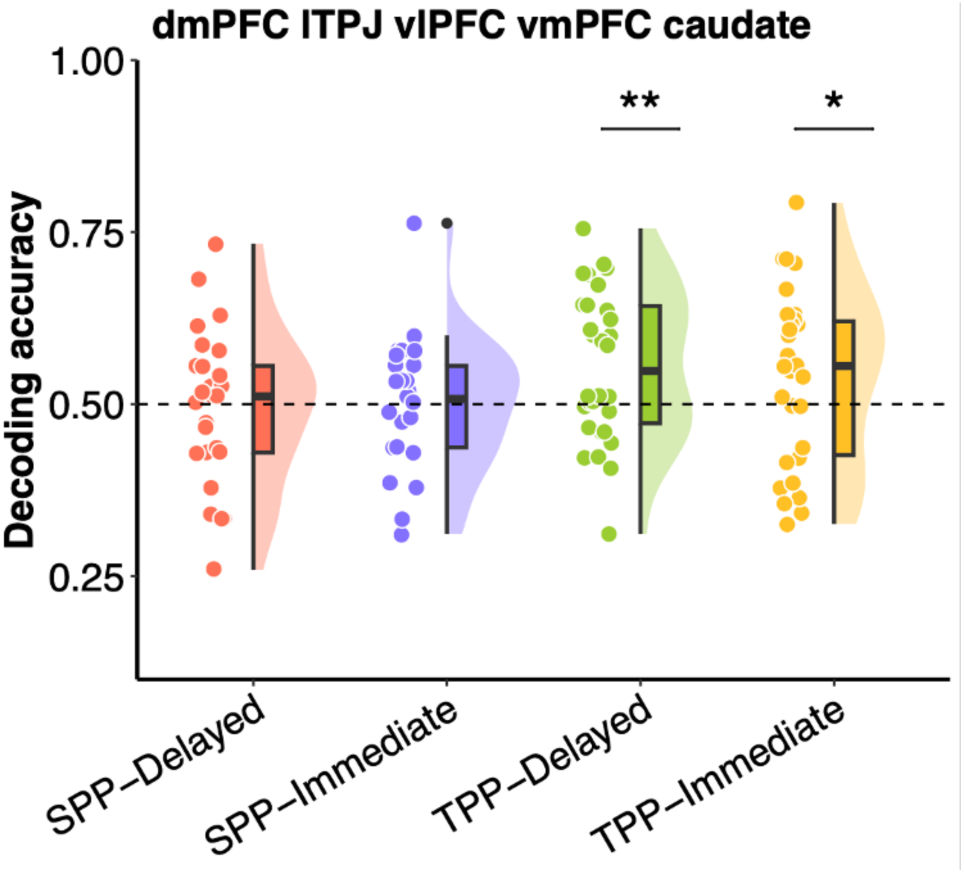
MVPA (multivariable pattern analysis) decoding results. When integrating dmPFC, TPJ, vlPFC, vmPFC, and caudate for model training, a significant prediction accuracy (> 0.5) was observed in the TPP delayed condition. * *p* <.05, ** *p* <.01.

#### Decoding Punishment Severity

We employed a MVPA (MVPV-Light toolbox, Treder, 2020) approach to examine the relationship between brain activity and punishment decision-making, utilizing beta values extracted from single-trial GLM to predict punishment levels. The GLM was constructed as the univariate analysis. Punishment levels were categorized as ‘high’ for ratings ≥ 5 and’low’ for ratings < 5. The rationale behind this approach is twofold: first, punishment severity is a central part of punishment decision-making. Brain regions capable of decoding punishment severity are expected to be closely involved in this processing. Furthermore, this approach advances the above univariate analyses to not only examine differences in punishment between immediate and delayed conditions but also to identify the brain regions closely associated with punishment severity. This allows for a deeper understanding of the neural mechanisms involved in these critical decision-making processes.

We adopted a ROI approach, selecting brain regions according to the HPM and relevant empirical research (see the Method section for details). A one-tailed *t*-test was performed to determine whether the prediction accuracy was significantly greater than 0.5 across SPP-immediate, SPP-delayed, TPP-immediate, and TPP-delayed conditions. Our findings revealed that when training the model using individual ROIs, no brain regions exhibited decoding accuracy significantly above chance, all *ps*_FDR_ >.05. Inspired by this finding, we hypothesized that the punishment decision-making might engage an integrative brain network rather than individual regions. To test this, we combined brain regions with significant activity differences between immediate and delayed conditions (i.e., the dmPFC, TPJ, vlPFC, vmPFC, and caudate) for model training. We observed significant prediction accuracy above chance in the TPP delayed condition (accuracy =.555, *t* (29) = 2.788, *p* =.0046), and TPP immediate condition (accuracy =.540, *t* (29) = 1.743, *p* =.046), with no such effect found in the other conditions. These results highlight the collaborative engagement of multiple brain regions in decision-making processes within the TPP contexts.

## Discussion

In real-world situations, the punishment of norm violators is often hindered as they flee, resulting in postponed consequences. However, little is known about how SPP and TPP, both vital for regulating social behavior and maintaining social norms, address these delayed sanctions. This study takes a departure from the conventional economic game paradigm (Civai et al., 2019; Stallen et al., 2018), and instead adopts real criminal scenario assessment approach. We have identified shared and distinct cognitive processes and neural substrates underlying immediate and delayed punishment in SPP and TPP, thereby contributing to the expanding field of neurolaw research.

The first objective of the present study was to examine the behavioral responses of SPP and TPP under varying temporal conditions. The findings suggested that delayed sanctions for transgressors elicited more severe punishment, stronger emotional arousal and a more negative life attitude when administered by a third-party. The more severe punishment may be attributed to two main explanations: first, the discounting of punishment in delayed sanctions, a phenomenon commonly observed in animal (Liley et al., 2019) and human studies (Bernasco et al., 2017; Madden et al., 2003); and second, the role of third parties as maintainers of social injustice, where delayed sanctions evoke a greater sense of unfairness compared to immediate sanction (Kundro et al., 2023). This aligns well with the stronger negative emotional arousal observed in the delayed condition, potentially leading to more severe punishment in the context of TPP. For SPP, participants exhibited stronger emotional arousal in the immediate sanctions compared to delayed sanction for the misdemeanor cases. These results indicate that participants focus more on the present rather than the delayed future, which may contribute to their irrational decision-making—assigning the same severity of punishment to both immediate and delayed sanctions. This study further identified the modulating effect of the severity of criminal scenarios. The delayed discounting effect of punishment primarily occurs in misdemeanor and felony cases, but not in capital offense cases. The disappearance of this effect in capital offense cases may be attributed to a ceiling effect, where the most severe punishment has already been assigned in both immediate and delayed contexts.

Regarding the neural correlates, we firstly compared the distinction between SPP and TPP. Starting from the HPM, we have the hypothesis that SPP may elicit stronger brain activity in brain regions associated with theory of mind than TPP. Our findings were consistent with this hypothesis, as we observed stronger brain activity in the precuneus and TPJ in SPP compared to TPP. Although not explicitly mentioned in HPM, the precuneus has also widely been identified as the functional core of DMN (Cavanna & Trimble, 2006; Utevsky et al., 2014). Both the precuneus and TPJ are consistently linked to perspective-taking and mentalizing, showing increased activity when individuals engage in tasks requiring them to adopt another person’s viewpoint and form representations of other’s mental states (Cavanna & Trimble, 2006; Feng et al., 2022, 2023; Saxe & Powell, 2006; Tholen et al., 2020; Yang et al., 2019; Young et al., 2007, 2010; Young & Saxe, 2008). The task-based FC analysis further revealed stronger connectivity between the left TPJ and the left dlPFC—a brain region closely associated with implementation of fairness motives (Buckholtz et al., 2015; Ginther et al., 2016; Knoch et al., 2006; Stallen et al., 2018)—in the SPP condition. Of note, these findings contrast with previous studies which usually report greater TPJ activity or TPJ-dlPFC connectivity (Bellucci et al., 2020; Feng et al., 2022; Yang et al., 2024) in TPP compared to SPP. These differences may stem from the experimental setup; In the current hypothetical criminal scenario paradigm, participants assumed the role of the victim in SPP or the role of the judge in TPP. Consequently, individuals in SPP group may perceive the hypothetical harm as more personally relevant compared to TPP group, leading them to engage more deeply with precuneus and TPJ to understand the perpetrator’s intentions, as well as enhancing connectivity between TPJ and dlPFC to arrive at a final decision.

When exploring the timing effect, whole brain analysis revealed stronger activation in the dmPFC during the delayed trial compared to the immediate trial. Importantly, this effect was observed exclusively in TPP, keeping consistent with the behavioral findings. The dmPFC is a component of the Theory of Mind (ToM) network, exhibiting heightened activation when individuals attempt to infer the thoughts of others (Isoda & Noritake, 2013; Krueger & Hoffman, 2016). Additionally, the dmPFC has been implicated in integrating contextual factors for making assessments of punishment (Bellucci et al., 2017; Feng et al., 2022; Ginther et al., 2016). This suggests that TPP individuals may uphold social norms by taking into account the reduced effectiveness of delayed sanctions and imposing harsher punishment on violators.

The ROIs analysis further revealed stronger activity in TPJ, vlPFC, vmPFC, and caudate during delayed sanctions compared to immediate sanctions in TPP. As mentioned earlier, the TPJ plays a key role in mentalizing (Saxe & Powell, 2006; Wu et al., 2020), i.e., simulating and understanding the perspectives of both the perpetrator and the victim to facilitate decision-making processes in the present study. The VLPFC, a key region involved in regulatory processes that promote fairness behaviors by inhibiting intuitive responses (Carlson & Crockett, 2018; Feng et al., 2015), has been found to be exclusively engaged in the TPP rather than SPP in one meta-analysis (Bellucci et al., 2020). The vmPFC and caudate are crucial for processing reward and punishment, as well as integrating emotional and social information to guide behavior (Bartra et al., 2013; Delgado, 2007; Koenigs & Tranel, 2007; Lo Gerfo et al., 2019; Pujara et al., 2016). Based on these findings, we deduced that a greater demand for information integration, value calculation, emotional regulation and executive control was required in the delayed sanctions compared to the immediate sanctions in TPP. Additionally, a stronger sense of reward was also elicited in the delayed sanctions, which can be attributed to the perception that maintaining social fairness through punishment over time is itself rewarding. This aligns with findings in Stallen et al., (2018). Similar to the behavioral findings, we also noted the modulating effect of the severity of the criminal scenario, with no significant difference between immediate and delayed sanctions in the capital offenses, further substantiating the ceiling effect in this condition.

The MVPA analysis further explored which brain regions were closely linked to punishment severity. The results indicated that using integrative ROIs, rather than individuals ROI for model training, led to higher decoding accuracy over chance in TPP delayed conditions, suggest a corroborative processing mechanism in punishment decision-making with the TPP context. These findings provide further support for the social brain network hypothesis (Blakemore, 2008), reinforcing the idea that multiple brain regions work together to facilitate behaviors and decision-making in social contexts.

The absence of significant differences in punishment assignment and brain regions between delayed and immediate sanctions in SPP suggests that the personal relevance (i.e., assuming being the victim vs. assuming being the judge) of harm may overshadow considerations of timing. It has been shown that episodic future thinking more effectively reduces monetary delay discounting when future events are self-relevant, as opposed to involving another person, highlight the role of self-relevance (Olsen et al., 2024). Our findings are consistent with this study and further extend this phenomenon to the field of delayed punishment. In SPP, where individuals were required to assume to be directly affected by the harm, the punishment of the perpetrator may prioritize addressing the harm they’ve experienced and consider less about the discounting utility of the delayed sanction, leading to comparable punishment severity and similar neural responses regardless of the timing of the sanction.

We also sought to investigate the association between brain activity and punishment ratings to identify the neural correlates of punishment discounting. Contrary to our expectation, we did not find significant correlations between brain activity difference (immediate vs. delayed) and punishment rating difference (immediate vs. delayed) in TPP group, where punishment discounting was present. However, we observed a significant negative correlation between activation differences and punishment differences in the TPJ and caudate in felony criminal scenarios within the SPP group. These findings suggest that greater self-engagement, as reflected by TPJ in SPP, was associated with smaller punishment differences between immediate and delayed sanctions, supporting the idea that the strong self-engagement in SPP diminishes the effect of sanction timing. Furthermore, the negative correlation between caudate activity and punishment severity difference indicates that the stronger the engagement of brain regions involved in emotional processing (Delgado, 2007), the more irrational punishment decision-making. In summary, these results enrich our understanding of the disappearance of delay discounting in SPP condition.

There were some limitations in the present study. Firstly, although we revealed how SPP and TPP process delayed sanctions, the emotional arousal ratings were collected after the fMRI scan, preventing us from thoroughly investigating the association between emotional arousal, punishment implementation and brain activities, as well as the potential role of emotional arousal in punishment decision-making, especially under the delayed condition. Future studies could address this by collecting emotional and behavioral ratings during the scan. Secondly, this study was unable to detect the gender difference in the delayed punishment due to the relatively small sample size. Liley et al. (2019) found that female rats discount delayed punishment less than males; However, researchers know little about sex differences in human sensitivity to delayed punishment discounting, along with the cognitive processes and neural systems underlying these differences. Future research with larger samples could explore this question and assess whether these findings generalize across species. Finally, a common yet noteworthy limitation is the selection of ROIs. Although these ROIs were chosen based on well-established studies, this approach does not encompass all possible brain regions, which may increase the risk of false negatives or false positives.

In summary, this was the first attempt to examine the behavioral and neural mechanisms underlying the decision-making process of second-and third-party punishment under time delay. Although participants in the current study were not a real judge or legal practitioner, the finding revealed how the general public understands and upholds fairness in the delayed sanction of perpetrators. The third-party perspective could shed light on the development of policy regarding delayed sentencing. Future studies may further explore this issue by employing legal practitioners.

### Methods Participants

We calculated the optimal sample size using G*Power 3.1.9.7 (Faul et al., 2007) for a medium effect size in a repeated measures design (*F*-test, with-between interaction, number of groups = 2, and number of measurements = 6) with the following parameters: groups = 2, *f* =.25, *α* =.05, 1 − *β* =.90, leading to a requirement of 24 participants in each group. Sixty-one Chinese proficient students, aged between 18 and 30 (mean age: 22.18 ± 3.18 years; 26 males), voluntarily enrolled in the study.

Participants were recruited through advertisements on university bulletin boards and social media platforms. Detailed demographic information is provided in Table S1. All participants were right-handed and had normal or corrected-to-normal vision. The exclusion criteria included participants with a history of psychiatric illness, those who had been victims of or witnesses to a violent crime (including sexual abuse), and those who had experienced any trauma involving injury or threat of injury, either as victims or as a close friend/family member of the victims. Participants were randomly assigned to SPP or TPP groups. 5 participants were excluded from the final analysis due to excessive head motion in the scanner. Consequently, the study included 26 participants in the SPP group (age: 22.15 ± 2.56 years; 9 males) and 30 participants in the TPP group (age: 22.30 ± 3.18 years; 14 males). Each participant provided written informed consent before the experiment and received monetary compensation of approximately MOP 150 for their participation. The study was approved by the Institutional Review Board of the University of Macau (BSERE21-APP005-ICI) and was conducted in strict accordance with the guidelines of research ethics.

### Experimental materials

Before the formal experiment, we collected 27 real legal cases (see SI Appendix for details) from the China Judgment Document Network (https://wenshu.court.gov.cn/) and recruited another cohort (41 participants, 23 females, ages: 19.951 ±1.161) online to assume the role of the second party (the victim) to provide initial ratings on criminal severity and emotional arousal for these cases. A law scholar in China has proposed that five years of fixed-term imprisonment should be the dividing line between misdemeanors and felonies in China’s criminal system (Bai, 2014). Using this criterion and the initial ratings, we divided the legal scenarios into three levels of seriousness, further distinguishing between immediate and delayed conditions.

Building on existing findings that stiffer criminal sanctions increase the avoidance efforts of offenders (Friehe & Miceli, 2017), we designed delay periods for criminal sentencing according to the severity of the crime. Specifically, for misdemeanors (N=9), where the defendants were sentenced to fines or short-term imprisonment for intentionally or negligently committing crimes that caused property losses and physical and mental injuries to victims, the delay times was set as the multiples of 0.1 years (0.1, 0.2, 0.3, 0.4, 0.5, 0.6, 0.7, 0.8, 0.9). For felony cases (N = 9), in which the defendants were sentenced to fixed-term imprisonment for intentionally or negligently committing crimes that caused severe damage to the victim’s property and physical and mental injuries, the delay times were multiples of 1 year (1, 2, 3, 4, 5, 6, 7, 8, 9). For capital offenses (N = 9), where the defendants were sentenced to death for deliberately committing crimes that resulted in the victim’s death, the delay times were expressed in multiples of 3 years (3, 6, 9, 12, 15, 18, 21, 24, 27). In total, 27 real legal cases written in Chinese were adapted. The word length of each case ranged from 92 to 190 words, with an average of 149.6 words.

### The criminal scenario assessment task

This experiment utilized a mixed design, incorporating a 2 (Time delay: immediate vs. delayed) × 2 (Group: SPP vs. TPP) × 3 (Criminal Scenario: misdemeanor vs. felony vs. capital offense) structure. Within this framework, time delay and criminal scenario were treated as a within-subject variables and group as a between-subject variable. Participants completed a modified version of the criminal scenario assessment task while under the fMRI scanner. In the instruction, participants were required to consider each legal case independently and were encouraged to use full scale for scoring. In the SPP group, participants assumed the role of the victim in a legal case, whereas in the TPP group, participants took the role of the judge.

The experiment proceeded as follows (Fig. 1): First, an infinite-time criminal scenario appeared on the screen. Below the scenario, ten gray rectangles marked with numbers (0, 1, 2, 3, 4, 5, 6, 7, 8, 9) remained visible until the participant pressed the mouse to determine the degree of punishment, ranging from 0 (no punishment) to 9 (extreme punishment). Participants made the punishment assessment twice according to different time delays: “If you are the victim (or the judge in the TPP group) of this case and the defendant was arrested by the police on the day of the crime (or N years after the crime in the delayed condition), how much punishment do you think the defendant should be given?” (see Fig. 1). After that, a small white square appeared for 6000-8000 ms, transitioning into a large white square that was displayed for 2000 ms before disappearing, indicating the end of the trial and the beginning of a new one. During the transition from the small to the large white square, participants were instructed to focus their eyes on the moment the small square transformed, without using the mouse to take any action. This task was designed to quickly shift their attention from the case to the expanding square, thereby minimizing the influence of the previous case’s punishment decision on the next case’s decision. In total, there were 54 trials, with 27 trials in both immediate and delayed punishment conditions. These legal cases were presented in a randomized sequence across participants using E-Prime 2.0 (Psychological Software Tools, Inc., Pittsburgh, PA) software.

After scanning, all participants completed emotional arousal ratings for each criminal scenario, using a scale from 0 (calm) *to* 9 (extremely excited). Participants in the TPP group were also required to complete an additional Life Attitude Questionnaire asking “How positive or negative do you feel about life?” with responses ranging from 0 (extremely positive) to 9 (extremely negative). Notably, participants in the SPP group were not asked to rate this due to the presence of deceased victims in some legal cases. Additionally, the Internal Scale Questionnaire (ISQ), which asked participants to imagine punishment level for scores of 1, 3, 5, 8, and 9 (Buckholtz et al., 2008), along with the Behavioral Inhibition System/Behavioral Activation System (BIS/BAS) Scale (Carver & White, 1994) were administered via an online questionnaire platform in mainland China (https://www.wjx.cn/). The results of these two questionnaires demonstrated a strong consistency among participants regarding their internal scale of justice and comparable sensitivity to punishment and reward between the two groups (see the SI Appendix Table S1).

### fMRI Data Acquisition

High-resolution anatomical and functional images were acquired using a Siemens Prisma 3T MRI scanner with a 32-channel head coil at the Center for Cognition and Brain Science, University of Macau. The visual display was presented on an LCD panel and back-projected onto a screen positioned at the front of the magnet bore. Participants lay supine in the scanner and viewed the display on a mirror positioned above them. Stimulus presentation was synchronized with fMRI volume acquisition, and participants’ responses were recorded using a fiber-optic mouse. For each participant, a high-resolution anatomical scan of the entire brain was collected using a T1-weighted 3D magnetization-prepared rapid acquisition with gradient echo (MP-RAGE) sequence with the following parameters: (1) repetition time (TR): 2300 ms; (2) echo time (TE): 2.26 ms; (3) inversion time (TI): 900 ms; (4) flip angle: 8°; (5) number of slices: 176; (6) Slice thickness: 1.0 mm; (7) field of view (FOV): 256 mm; (8) matrix size: 256 × 256; (9) voxel size: 1 × 1 × 1 mm^3^. The blood oxygen level-dependent (BOLD) signal for functional images was measured using a T2-weighted gradient-echo planar imaging (EPI) sequence with the following parameters: (1) repetition time (TR): 1000 ms; (2) echo time (TE): 30.0 ms; (3) flip angle: 90°; (4) number of slices: 65; (5) Slice thickness: 2.0 mm; (6) field of view (FOV): 192 mm; (7) matrix size: 96×96; (8) voxel size: 2 × 2 × 2 mm^3^.

### Behavioral data analysis

All statistical analyses of the behavioral data were conducted with SPSS 20 for PC (SPSS Inc., Chicago, IL, USA). 2 (Time delay: immediate vs. delayed) × 3 (Criminal scenario: misdemeanor vs. felony vs. capital offense) × 2 (Group: SPP vs. TPP) three-way mixed designed ANOVA was conducted on the degree of the punishment and emotional arousal. Additionally, a 2 (Time delay: immediate vs. delayed) × 3 (Criminal scenario: misdemeanor vs. felony vs. capital offense) two-way repeated measures ANOVA was carried out on attitudes to life. If the sphericity assumption was violated according to the Mauchly’s test, Greenhouse-Geisser correction was applied to adjust the degrees of freedom for the *F*-tests of within-subject effects. The results were visualized using R 4.4.0.

### fMRI data analysis Preprocessing

fMRI data were preprocessed with Data Processing Assistant for fMRI Advanced Edition (DPARSF V5.0) (Yan et al., 2016), a MATLAB-based package built on the Statistical Parametric Mapping platform (SPM12). Specifically, slice timing was performed to align the scanning time delay for each slice within a single TR, followed by realignment to reduce the impact of head motion. The images were then co-registered to the participant’s anatomical image. To enable inter-subject comparison in a standard space, each individual’s functional images were transformed to MNI space using the conversion algorithms in the SPM package. Additionally, the functional neuroimaging data were smoothed with a 4 mm kernel to increase statistical robustness.

### Univariate analysis

#### Whole brain analyses

After the preprocessing, the data from all sessions were concatenated together. To extract activated brain regions associated with the processing of six different conditions**—**including three categories of severity at two types of time points**—**we applied general linear modeling (GLM) to probe the manipulative experimental effects on the brain region hemodynamics for each participant. In each GLM model, the regressors included the indicators for the six manipulative conditions (2 (Time delay: immediate vs. delayed) × 3 (Criminal Scenario: misdemeanor vs. felony vs. capital offense)) and six head-motion parameters. These regressors were convolved with the canonical hemodynamic response function (HRF) and its time derivatives.

The contrasts extracted from the within-subject level were used to perform between-subject statistical analyses in the second-level analyses. We first compared neural differences between the SPP and TPP groups. To investigate the effect of time delay, we performed a paired sample *t*-test for the time delay condition (delayed vs. immediate) separately in the SPP and TPP groups. All results presented in the second-level analysis were corrected using the family-wise error (FWE) method to reduce multiple comparison bias (*p* <.001 uncorrected, FWE cluster-level corrected *q* <.05). Additionally, to minimize false positive rates, non-parametric permutation testing (5000 times) (Eklund et al., 2016) was applied. The results of the fMRI data were then visualized using MRIcron (https://www.nitrc.org/projects/mricron), with brain regions marked using coordinates according to the Automatic Anatomical Labeling (AAL) template.

### Region of interests (ROI) analyses

ROI analysis was conducted within the framework of the HPM to identify the neural differences between the groups (SPP vs. TPP) and time delays (immediate vs. delayed). For the SPP vs. TPP comparison, the TPJ was selected as an ROI based on significant activation of the precuneus (peaked in the left hemisphere) observed in the whole-brain analysis, as well as its central role in theory of mind. For the comparison between delayed and immediate sanctions, key brain regions of DMN, including left TPJ, and vmPFC, were chosen as ROIs, given their involvement in information integration (Bellucci et al., 2020; Krueger & Hoffman, 2016). Additionally, the vlPFC and caudate were selected as ROIs based on findings from a meta-analysis study of social punishment, which reported stronger activation in the left vlPFC during TPP (Bellucci et al., 2020), as well as the hypothesis that upholding social norms by punishing norm violators is perceived as rewarding, particularly in the delayed sanctions (de Quervain et al., 2004; Stallen et al., 2018; Strobel et al., 2011).

Among the selected ROIs, vmPFC (MNI: x/y/z = 2/54/– 4) (de Quervain et al., 2004), VLPFC(MNI: x /y/z =-46/30/-16) (Bellucci et al., 2020), caudate (MNI: x/y/z = 10/10/-2) (Strobel et al., 2011) were defined as 8-mm-radius spheres using by MarsBaR (Brett et al., 2002). TPJ (MNI: x/y/z =-54/-46/6) was retrieved from http://www.rbmars.dds.nl/CBPatlases.htm. The ROI analysis involved extracting the beta value from each ROI, followed by independent *t*-tests to compare group differences and paired *t*-tests to assess the effect of delay effect. The false discovery rate (FDR, *p* <.05) was applied to correct for multiple comparison.

### Task-based functional connectivity (FC)

Previous studies have identified a key role of the dlPFC in punishment implementation (Buckholtz et al., 2015; Knoch et al., 2006). Feng et al. (2022) revealed a stronger FC between right TPJ and right dlPFC during TPP compared to SPP. Integrating these findings and our observation that stronger activation in left TPJ was elicited in SPP compared to TPP, we conducted a task-based FC analysis between the left TPJ and left dlPFC to investigate the FC differences between SPP and TPP (Rissman et al., 2004). Specifically, we extracted the average time series for each ROI separately and calculated the Pearson correlations between ROIs for each participant.

### Correlation of brain activation and punishment ratings in immediate vs. delayed conditions

We investigated the relationship between brain activity differences and behavior ratings differences across immediate and delayed conditions to determine whether and which brain regions activity could explain the punishment-discounting effect. To do this, we selected regions that exhibited significant differences between immediate and delayed sanctions as ROIs. In the present study, they were left dmPFC, right vmPFC, left vlPFC, right TPJ, and right caudate (See Results section for details). We used the neural differences (immediate vs. delayed) in these regions as a neural index and the difference in punishment ratings between these conditions as a behavioral measure of punishment discounting. Pearson correlation analysis was then conducted to determine whether brain activation could explain the punishment-discounting effect.

### Multivariate pattern analysis (MVPA)

To further investigate the association between brain activity and punishment implementation, a MVPA approach was adopted. Initially, a general linear model (GLM) analysis was conducted for each trial involving each participant. Different with the univariate analysis, the dependent variable was the unsmoothed functional images. Next, a ROI approach was employed to extract beta values from specific brain regions that showed significant activation differences between immediate and delayed sanctions. We selected the brain regions under HPM as ROIs, including dmPFC, TPJ, vmPFC, vlPFC and Caudate (see the ROI analyses for details). These beta values were then utilized to predict punishment levels, defined as ‘high’ for ratings ≥ 5 and ‘low’ for ratings < 5, using a machine learning algorithm. Specifically, for each participant, the mv_classify function from the MVPV-Light toolbox (Treder, 2020) was invoked to classify the punishment levels based on the extracted beta values. We conducted this analysis across all cases rather than by severity, as punishment ratings in misdemeanor and capital offense cases may consistently be above 5 or below 5.

To assess the model’s predictive performance, linear discriminant analysis (LDA) was employed as the classifier, incorporating five-fold cross-validation (k-fold=5). The dataset was divided into five subsets, or folds, with four folds used to train the LDA model, allowing it to learn patterns associated with punishment severity. The remaining fold served as the test set to evaluate the model’s performance on unseen data, yielding performance metrics for each participant. Subsequently, *t*-test (one-tail) was performed to determine whether the prediction accuracy was significantly greater than 0.5 across the four conditions: SPP immediate, SPP delayed, TPP immediate, and TPP delayed. Through this method, we aimed to identify neural patterns associated with punishment decision-making and explore the influence of current versus delayed conditions on individuals’ punitive behavior.

## Data, Materials, and Software Availability

Codes reproducing the main figures can be found at GitHub (https://github.com/andlab-um/neuroLaw). All criminal cases description in the experiment are included in the manuscript and/or SI Appendix.

## Acknowledgments

This work was supported by FDCT of Macau [0127/2020/A3, 0041/2022/A], the Natural Science Foundation of Guangdong Province (2021A1515012509), Shenzhen-Hong Kong-Macao Science and Technology Innovation Project (Category C) (SGDX2020110309280100), MYRG of University of Macau (MYRG2022-00188-ICI), Humanities and Social Sciences Research Project of the Ministry of Education (24YJA190011), and Henan Province Key Scientific Research Project Plan for Higher Education Institutions (25A190002).

