## Supplemental figures, tables for "Unraveling the Neurocognitive Mechanisms of Delayed Punishment in Second-and Third-Party Contexts"

### Supplemental materials

#### Figures

Figure S1

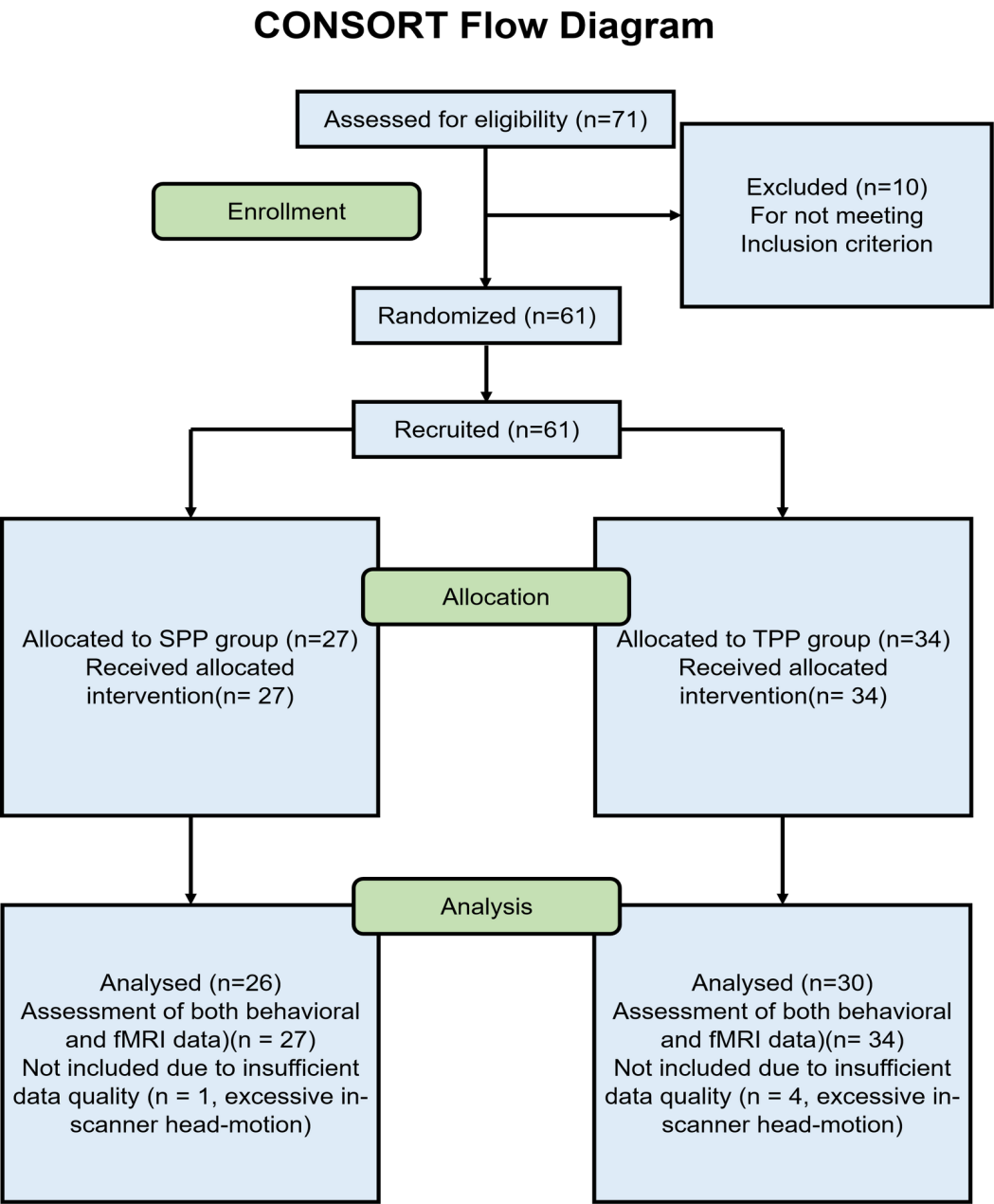

Figure S1: The CONSORT flow diagram of the study. All participant was recruited through advertisement. Six participants were excluded for not meeting the inclusion criterion. After the screening of the inclusion criterion, 60 participants were asked to have a lab visit and perform the (SPP/TPP) treatment randomly and finish the fMRI scanning. After that, we performed the questionnaire follow-up sessions, including the side effect checking and mood state changes. In the follow-up fMRI data analysis, one participant of the SPP group was excluded due to insufficient data quality (n=1, excessive in-scanner head-motion with motion criteria (> 3 mm translation or > 3 degree rotation)

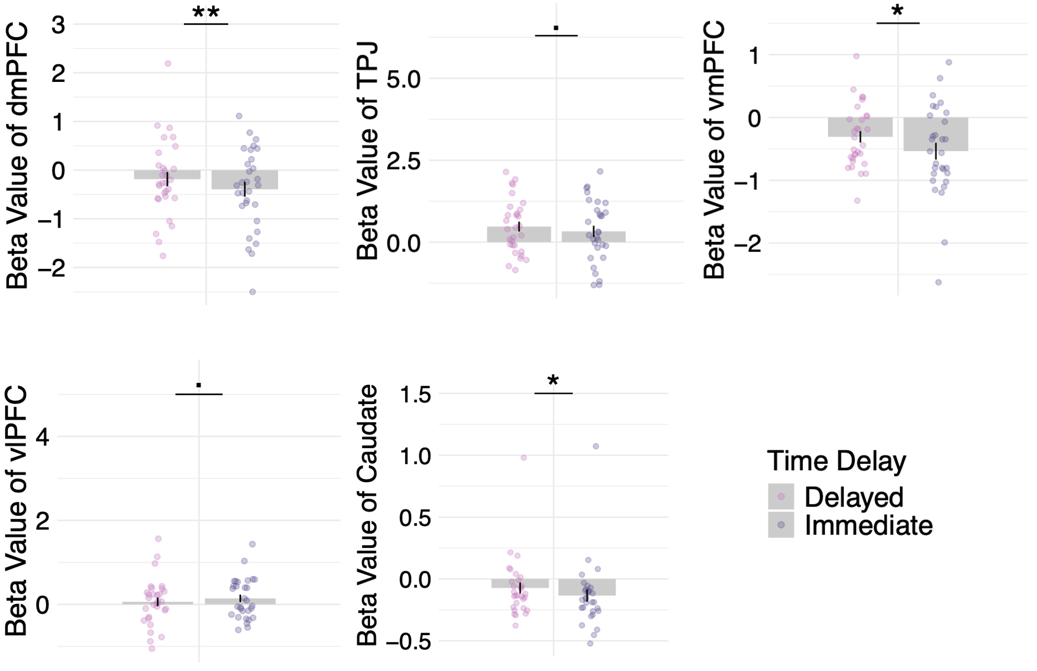

Figure S2. Differences in brain responses between immediate and delayed sanctions in the third-party punishment (TPP) group in felony cases. ROI analyses showed that the activities of dorsomedial prefrontal cortex (dmPFC, MNI: x/y/z = -14/60/26), temporoparietal junction (TPJ, MNI: x/y/z = -54/-46/6), ventromedial prefrontal cortex (vmPFC, MNI: x/y/z = 2/54/-4), ventrolateral prefrontal cortex (vlPFC, MNI: x/y/z = -46/30/-16) and caudate (MNI: x/y/z = 10/10/-2) were stronger in the delayed condition compared to the immediate condition in the TPP group during felony condition. *p* < 0.1, * *p* < .05, *p* < 0.01.

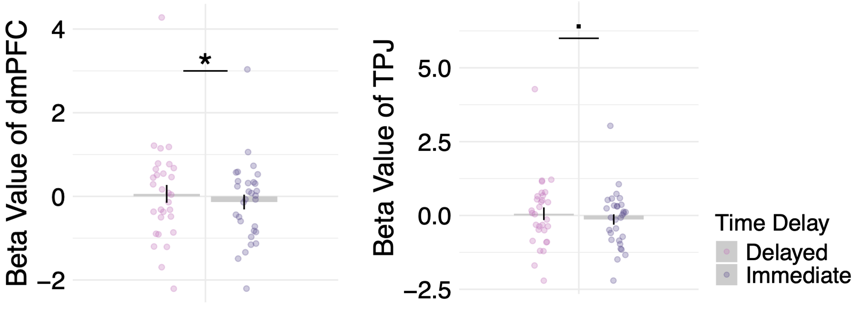

Figure S3. Differences in brain responses between immediate and delayed sanctions in the third-party punishment (TPP) group in misdemeanor cases. ROI analyses showed that activities in the dorsomedial prefrontal cortex (dmPFC, MNI: x/y/z = -14/60/26) and temporoparietal junction (TPJ, MNI: x/y/z = -54/-46/6) were stronger in the delayed condition compared to the immediate condition in the TPP group during felony cases. *p* < 0.1 * *p* < .05

Tables

Table S1. Participants’ demographic information

|  | SPP | TPP | *t*-score/Chi-square | *p*-value |
| --- | --- | --- | --- | --- |
| Age | 22.08 (2.637) | 22.30 (3.239) | -0.280 | 0.781 |
| Gender | 10 females | 16 females | 0.254 | 0.614 |
| Internal Scale Questionnaire | 1 = Monetary Fine  (84.6%)  3 = Detention or short-term imprisonment  (96.2%)  5 = Medium- to long-term imprisonment  (57.7%)  8 = Long-term imprisonment to life imprisonment  (100.0%)  9 = Death penalty  (96.1%) | 1 = Monetary Fine  (93.3%)  3 = Detention or short-term imprisonment  (96.7%)  5 = Medium- to long-term imprisonment  (50.0%)  8 = Long-term imprisonment to life imprisonment  (100.0%)  9 = Death penalty  (86.7%) | A strong consistency |  |
| BAS | 23.12 (5.094) | 23.60 (5.243) | -0.350 | 0.728 |
| BIS | 8.92 (2.576) | 10.47 (3.540) | -1.840 | 0.071 |

*Note*: Means (SD). BAS: Behavioral activation system, BIS, behavioral inhibition system.

Table S2. The brain activity in SPP and TPP

| Brain regions | | | MNI coordinates | | | Max *F*-value | *p*  (FWE-corr.) | Voxel size |
| --- | --- | --- | --- | --- | --- | --- | --- | --- |
|  |  |  | x | y | z |  |  |  |
| SPP group | | |  |  |  |  |  |  |
|  | Middle occipital gyrus _L | | -34 | -66 | 20 | 40.229 | 0.000 | 3331 |
|  | Middle temporal gyrus _R | | 52 | -46 | -8 | 29.01 | 0.000 | 6102 |
|  | Insular _R | | 30 | 28 | 4 | 17.922 | 0.000 | 411 |
|  | Inferior frontal gyrus _L | | -50 | 30 | 12 | 18.410 | 0.000 | 180 |
|  | Middle frontal cortex _R | | 32 | 4 | 42 | 13.465 | 0.028 | 70 |
|  | Precentral gyrus _L | | -40 | -2 | 42 | 12.212 | 0.003 | 112 |
| TPP group | | |  |  |  |  |  |  |
|  | | Middle temporal gyrus _L | -52 | -52 | 22 | 36.272 | 0.000 | 6022 |
|  | | Rolandic operculum _R | 46 | -30 | -14 | 50.256 | 0.000 | 12480 |
|  | | Inferior frontal gyrus _L | -50 | 26 | -14 | 24.456 | 0.000 | 2243 |
|  | | Precuneus _R | 30 | -52 | -2 | 21.231 | 0.000 | 332 |
|  | | Anterior cingulate cortex _R | 2 | 34 | -2 | 19.096 | 0.000 | 553 |
|  | | Occipital lobe _R | 26 | -64 | 32 | 12.382 | 0.014 | 97 |
|  | | Medial and paracingulate gyrus _R | 4 | -36 | 34 | 12.340 | 0.045 | 75 |
|  | | Superior frontal gyrus _L | -8 | 46 | 48 | 13.771 | 0.005 | 119 |
| Across groups | | |  |  |  |  |  |  |
|  | | Superior temporal gyrus _L | -54 | -2 | 24 | 14.952 | 0.000 | 155 |
|  | | Inferior temporal gyrus _L | -42 | -48 | -14 | 15.143 | 0.000 | 138 |
|  | | Lingual gyrus _L | -8 | -68 | -6 | 35.428 | 0.001 | 418 |
|  | | Lingual gyrus _R | 8 | -68 | 0 | 27.950 | 0.001 | 77 |
|  | | Lingual gyrus _R | 20 | -62 | 0 | 19.528 | 0.000 | 112 |
|  | | Middle temporal gyrus _L | -60 | -54 | 20 | 13.693 | 0.000 | 209 |
|  | | Inferior parietal lobule _R | 42 | -54 | 39 | 13.876 | 0.000 | 148 |
|  | | Supramarginal gyrus _R | 48 | -32 | 46 | 16.781 | 0.000 | 89 |
|  | | Precuneus _R | 10 | -56 | 50 | 18.710 | 0.001 | 202 |
|  | | Middle frontal gyrus _R | 26 | 16 | 48 | 14.636 | 0.001 | 83 |
|  | | Precuneus _R | 10 | -66 | 52 | 17.208 | 0.000 | 142 |

Table S3. The brain activity of timing delay effect

| Brain regions | MNI coordinates | | | Max *t*-value | *p* (FWE-corr.) | Voxel size | |
| --- | --- | --- | --- | --- | --- | --- | --- |
|  | x | y | z |  |  |  |  |
| SPP: Immediate - Delayed |  |  |  |  |  | |  |
| No significant cluster |  |  |  |  |  | |  |
| SPP: Delayed -Immediate |  |  |  |  |  | |  |
| No significant cluster |  |  |  |  |  | |  |
| TPP: Immediate - Delayed |  |  |  |  |  | |  |
| No significant cluster |  |  |  |  |  | |  |
| TPP: Delayed -Immediate |  |  |  |  |  | |  |
| Dorsolateral superior frontal gyrus _L | -14 | 60 | 26 | 4.681 | 0.018 | | 91 |
| Across groups: Immediate - Delayed |  |  |  |  |  | |  |
| no significant cluster |  |  |  |  |  | |  |
| Across groups: Delayed -Immediate |  |  |  |  |  | |  |
| Dorsolateral superior frontal gyrus _L | -20 | 8 | -6 | 4.790 | 0.099 | | 57 |

Experimental materials

The criminal cases used in the experiment were all sourced from China Judgment Document Network (<https://wenshu.court.gov.cn/>).

Misdemeanor

(1) On December 3, 2012, the victim, Liu, purchased goods at a supermarket in Suzhou City using cash. Due to an urgent matter, Liu left the store without counting the change. Later, upon counting the change, Liu discovered that the supermarket was short by several dozen yuan and concluded that the supermarket owner (defendant Zhang) had intentionally shortchanged him. Liu subsequently reported the incident to the police.

Second Party

1. If you were the victim in this case and the defendant was arrested by the police on the day of the crime, what level of punishment do you believe should be imposed on the defendant?

2. If you were the victim in this case and the defendant was arrested by the police on the day of the crime, what would be your level of emotional distress?

3. If you were the victim in this case and the defendant was arrested by the police one month later, what level of punishment do you believe should be imposed on the defendant?

4. If you were the victim in this case and the defendant was arrested by the police one month later, what would be your level of emotional distress?

Third Party

5. If you were the judge in this case and the defendant was arrested by the police on the day of the crime, what level of punishment do you believe should be imposed on the defendant?

6. If you were the judge in this case and the defendant was arrested by the police on the day of the crime, what would be your attitude towards the defendant's sentence?

7. If you were the judge in this case and the defendant was arrested by the police on the day of the crime, what would be your level of emotional distress?

8. If you were the judge in this case and the defendant was arrested by the police one month later, what level of punishment do you believe should be imposed on the defendant?

9. If you were the judge in this case and the defendant was arrested by the police one month later, what would be your attitude towards the defendant's sentence?

10. If you were the judge in this case and the defendant was arrested by the police one month later, what would be your level of emotional distress?

(2) On October 13, 2018, the defendant, Shu, stole a pair of pants and two small scissors with a total value exceeding 100 yuan from a store. The store manager, Li, was unaware of the theft at the time. Upon counting the inventory, Li discovered discrepancies and reviewed the surveillance footage, which revealed that someone had stolen goods from the supermarket. Li then reported the incident to the police.

Second Party

1. If you were the victim in this case and the defendant was arrested by the police on the day of the crime, what level of punishment do you believe should be imposed on the defendant?

2. If you were the victim in this case and the defendant was arrested by the police on the day of the crime, what would be your level of emotional distress?

3. If you were the victim in this case and the defendant was arrested by the police two months later, what level of punishment do you believe should be imposed on the defendant?

4. If you were the victim in this case and the defendant was arrested by the police two months later, what would be your level of emotional distress?

Third Party

5. If you were the judge in this case and the defendant was arrested by the police on the day of the crime, what level of punishment do you believe should be imposed on the defendant?

6. If you were the judge in this case and the defendant was arrested by the police on the day of the crime, what would be your attitude towards the defendant's sentence?

7. If you were the judge in this case and the defendant was arrested by the police on the day of the crime, what would be your level of emotional distress?

8. If you were the judge in this case and the defendant was arrested by the police two months later, what level of punishment do you believe should be imposed on the defendant?

9. If you were the judge in this case and the defendant was arrested by the police two months later, what would be your attitude towards the defendant's sentence?

10. If you were the judge in this case and the defendant was arrested by the police two months later, what would be your level of emotional distress?

(3) On January 21, 2018, the defendant, Peng, used self-media accounts registered on various platforms (such as Weibo) to publish articles about the private lives of entertainers, specifically targeting the victim, Yang. The articles were filled with mocking and derogatory language, and continued to "criticize" the victim, Yang. This objectively led readers to lower their evaluation of the victim's character, thereby constituting an infringement on the victim's reputation. The victim, Yang, subsequently filed a lawsuit with the court.

Second Party

1. If you were the victim in this case and the defendant was arrested by the police on the day of the crime, what level of punishment do you believe should be imposed on the defendant?

2. If you were the victim in this case and the defendant was arrested by the police on the day of the crime, what would be your level of emotional distress?

3. If you were the victim in this case and the defendant was arrested by the police nine months later, what level of punishment do you believe should be imposed on the defendant?

4. If you were the victim in this case and the defendant was arrested by the police nine months later, what would be your level of emotional distress?

Third Party

5. If you were the judge in this case and the defendant was arrested by the police on the day of the crime, what level of punishment do you believe should be imposed on the defendant?

6. If you were the judge in this case and the defendant was arrested by the police on the day of the crime, what would be your attitude towards the defendant's sentence?

7. If you were the judge in this case and the defendant was arrested by the police on the day of the crime, what would be your level of emotional distress?

8. If you were the judge in this case and the defendant was arrested by the police nine months later, what level of punishment do you believe should be imposed on the defendant?

9. If you were the judge in this case and the defendant was arrested by the police nine months later, what would be your attitude towards the defendant's sentence?

10. If you were the judge in this case and the defendant was arrested by the police nine months later, what would be your level of emotional distress?

(4) On June 2, 2016, the victim, Kuang, was driving a bus to a bus stop when he noticed two broken coins in the defendant, Li's, hand. Kuang requested that Li exchange the coins, and Li proceeded to trade money with other passengers. Kuang continued to drive the bus, but Li repeatedly insulted him during the journey. When Li struck Kuang's head, Kuang stopped the bus and confronted him. The defendant then struck the coin machine while grabbing Kuang's head. A passenger dialed 911, and the defendant subsequently jumped out of the window. The victim's injuries were determined to be at disability level 10 following identification.

Second Party

1. If you were the victim in this case and the defendant was arrested by the police on the day of the crime, what level of punishment do you believe should be imposed on the defendant?

2. If you were the victim in this case and the defendant was arrested by the police on the day of the crime, what would be your level of emotional distress?

3. If you were the victim in this case and the defendant was arrested by the police six months later, what level of punishment do you believe should be imposed on the defendant?

4. If you were the victim in this case and the defendant was arrested by the police six months later, what would be your level of emotional distress?

Third Party

5. If you were the judge in this case and the defendant was arrested by the police on the day of the crime, what level of punishment do you believe should be imposed on the defendant?

6. If you were the judge in this case and the defendant was arrested by the police on the day of the crime, what would be your attitude towards the defendant's sentence?

7. If you were the judge in this case and the defendant was arrested by the police on the day of the crime, what would be your level of emotional distress?

8. If you were the judge in this case and the defendant was arrested by the police six months later, what level of punishment do you believe should be imposed on the defendant?

9. If you were the judge in this case and the defendant was arrested by the police six months later, what would be your attitude towards the defendant's sentence?

10. If you were the judge in this case and the defendant was arrested by the police six months later, what would be your level of emotional distress?

(5) On August 9, 2018, the defendant, Cui, was driving a small cross-country bus toward a highway entrance when he failed to yield to a vehicle traveling straight ahead, causing a collision between Cui's vehicle and the victim, Zhang's minivan. This resulted in damage to both vehicles and injuries to Zhang. After the accident, Cui abandoned the vehicle and fled the scene. The victim, Zhang, was admitted to the hospital for medical treatment. Following an assessment, Zhang's injuries were determined to be at disability level 10. The traffic police concluded that the defendant, Cui, fled the scene after abandoning the vehicle and was held fully responsible for the collision.

Second Party

1. If you were the victim in this case and the defendant was arrested by the police on the day of the crime, what level of punishment do you believe should be imposed on the defendant?

2. If you were the victim in this case and the defendant was arrested by the police on the day of the crime, what would be your level of emotional distress?

3. If you were the victim in this case and the defendant was arrested by the police eight months later, what level of punishment do you believe should be imposed on the defendant?

4. If you were the victim in this case and the defendant was arrested by the police eight months later, what would be your level of emotional distress?

Third Party

5. If you were the judge in this case and the defendant was arrested by the police on the day of the crime, what level of punishment do you believe should be imposed on the defendant?

6. If you were the judge in this case and the defendant was arrested by the police on the day of the crime, what would be your attitude towards the defendant's sentence?

7. If you were the judge in this case and the defendant was arrested by the police on the day of the crime, what would be your level of emotional distress?

8. If you were the judge in this case and the defendant was arrested by the police eight months later, what level of punishment do you believe should be imposed on the defendant?

9. If you were the judge in this case and the defendant was arrested by the police eight months later, what would be your attitude towards the defendant's sentence?

10. If you were the judge in this case and the defendant was arrested by the police eight months later, what would be your level of emotional distress?

(6) On February 23, 2020, the victim, Gong, was walking down a road in a village in Yingshan County when Zhou's family cow, which was grazing in the field without a rope, heard the noise of a passing motorcyclist and ran toward the road. Gong was knocked down and injured while trying to avoid the cow. He was subsequently taken to the hospital for medical treatment. Gong's family then called the police. Following an assessment, Gong's injuries were determined to be at disability level 10.

Second Party

1. If you were the victim in this case and the defendant was arrested by the police on the day of the crime, what level of punishment do you believe should be imposed on the defendant?

2. If you were the victim in this case and the defendant was arrested by the police on the day of the crime, what would be your level of emotional distress?

3. If you were the victim in this case and the defendant was arrested by the police four months later, what level of punishment do you believe should be imposed on the defendant?

4. If you were the victim in this case and the defendant was arrested by the police four months later, what would be your level of emotional distress?

Third Party

5. If you were the judge in this case and the defendant was arrested by the police on the day of the crime, what level of punishment do you believe should be imposed on the defendant?

6. If you were the judge in this case and the defendant was arrested by the police on the day of the crime, what would be your attitude towards the defendant's sentence?

7. If you were the judge in this case and the defendant was arrested by the police on the day of the crime, what would be your level of emotional distress?

8. If you were the judge in this case and the defendant was arrested by the police four months later, what level of punishment do you believe should be imposed on the defendant?

9. If you were the judge in this case and the defendant was arrested by the police four months later, what would be your attitude towards the defendant's sentence?

10. If you were the judge in this case and the defendant was arrested by the police four months later, what would be your level of emotional distress?

(7) On March 16, 2016, the basketball team of the defendant, Guo, lost to its opponent in the finals. After the game, Guo returned to the hotel where he was staying, and a fight broke out between the fans of Guo's team and the opposing team's fans. Witnessing his parents being attacked by the opposing fans, Guo intervened to protect his family and physically assaulted the victim, Zeng. Zeng's injuries were classified as a Grade 10 disability. Based on the results of the investigation and treatment by the public security authorities, and in accordance with the disciplinary provisions of the league, the Basketball Association took action against the defendant, Guo.

Second Party

1. If you were the victim in this case and the defendant was arrested by the police on the day of the crime, what level of punishment do you believe should be imposed on the defendant?

2. If you were the victim in this case and the defendant was arrested by the police on the day of the crime, what would be your level of emotional distress?

3. If you were the victim in this case and the defendant was arrested by the police five months later, what level of punishment do you believe should be imposed on the defendant?

4. If you were the victim in this case and the defendant was arrested by the police five months later, what would be your level of emotional distress?

Third Party

5. If you were the judge in this case and the defendant was arrested by the police on the day of the crime, what level of punishment do you believe should be imposed on the defendant?

6. If you were the judge in this case and the defendant was arrested by the police on the day of the crime, what would be your attitude towards the defendant's sentence?

7. If you were the judge in this case and the defendant was arrested by the police on the day of the crime, what would be your level of emotional distress?

8. If you were the judge in this case and the defendant was arrested by the police five months later, what level of punishment do you believe should be imposed on the defendant?

9. If you were the judge in this case and the defendant was arrested by the police five months later, what would be your attitude towards the defendant's sentence?

10. If you were the judge in this case and the defendant was arrested by the police five months later, what would be your level of emotional distress?

(8) On July 6, 2018, the victim, Zhao, purchased Japanese health products from the Taobao store operated by the defendant, Zhong, for a total of approximately 3,000 yuan. These products were sold without the necessary product sales number or drug registration number. They were imported from Japan and lacked Chinese labeling. Additionally, they did not comply with the requirements of the national quarantine and entry-exit inspection department for certification materials. According to the law, these products were deemed to be food items that did not meet food safety standards. Zhao filed a lawsuit with the court, alleging that the products did not have the effects claimed by Zhong.

Second Party

1. If you were the victim in this case and the defendant was arrested by the police on the day of the crime, what level of punishment do you believe should be imposed on the defendant?

2. If you were the victim in this case and the defendant was arrested by the police on the day of the crime, what would be your level of emotional distress?

3. If you were the victim in this case and the defendant was arrested by the police seven months later, what level of punishment do you believe should be imposed on the defendant?

4. If you were the victim in this case and the defendant was arrested by the police seven months later, what would be your level of emotional distress?

Third Party

5. If you were the judge in this case and the defendant was arrested by the police on the day of the crime, what level of punishment do you believe should be imposed on the defendant?

6. If you were the judge in this case and the defendant was arrested by the police on the day of the crime, what would be your attitude towards the defendant's sentence?

7. If you were the judge in this case and the defendant was arrested by the police on the day of the crime, what would be your level of emotional distress?

8. If you were the judge in this case and the defendant was arrested by the police seven months later, what level of punishment do you believe should be imposed on the defendant?

9. If you were the judge in this case and the defendant was arrested by the police seven months later, what would be your attitude towards the defendant's sentence?

10. If you were the judge in this case and the defendant was arrested by the police seven months later, what would be your level of emotional distress?

(9) On May 6, 2015, the victim, Wang, took her granddaughter to the supermarket, where the defendant, Liu, served as the legal representative. Wang's granddaughter pushed a baby stroller and walked up the escalator on her own. The supermarket staff did not intervene to stop her granddaughter's behavior, and she fell backward after standing unsteadily on the escalator. As a public place, the supermarket has a duty to protect the personal safety of its patrons within a reasonable limit, but the supermarket staff failed to stop the escalator in time to prevent the fall. Wang subsequently filed a lawsuit against the defendant, Liu.

Second Party

1. If you were the victim in this case and the defendant was arrested by the police on the day of the crime, what level of punishment do you believe should be imposed on the defendant?

2. If you were the victim in this case and the defendant was arrested by the police on the day of the crime, what would be your level of emotional distress?

3. If you were the victim in this case and the defendant was arrested by the police three months later, what level of punishment do you believe should be imposed on the defendant?

4. If you were the victim in this case and the defendant was arrested by the police three months later, what would be your level of emotional distress?

Third Party

5. If you were the judge in this case and the defendant was arrested by the police on the day of the crime, what level of punishment do you believe should be imposed on the defendant?

6. If you were the judge in this case and the defendant was arrested by the police on the day of the crime, what would be your attitude towards the defendant's sentence?

7. If you were the judge in this case and the defendant was arrested by the police on the day of the crime, what would be your level of emotional distress?

8. If you were the judge in this case and the defendant was arrested by the police three months later, what level of punishment do you believe should be imposed on the defendant?

9. If you were the judge in this case and the defendant was arrested by the police three months later, what would be your attitude towards the defendant's sentence?

10. If you were the judge in this case and the defendant was arrested by the police three months later, what would be your level of emotional distress?

Felony

(1) On June 10, 2018, the defendant, Nan's husband, was arrested by the police station on suspicion of drug abuse. The following day, in an attempt to secure her husband's release, the defendant, Nan, pretended to report drug users but instead lured others into drug use and then brought them to the Public Security Bureau to surrender, hoping to create meritorious deeds for her husband. Nan contacted the victim, Chen, and arranged for him to use drugs at a hotel. After Chen's drug use, Nan took him to a designated location and called the police. Chen was subsequently arrested by the police and received an administrative penalty. In return, Nan paid Chen a remuneration of 1,000 yuan.

Second Party

1. If you were the victim in this case and the defendant was arrested by the police on the day of the crime, what level of punishment do you believe should be imposed on the defendant?

2. If you were the victim in this case and the defendant was arrested by the police on the day of the crime, what would be your level of emotional distress?

3. If you were the victim in this case and the defendant was arrested by the police one year after the crime, what level of punishment do you believe should be imposed on the defendant?

4. If you were the victim in this case and the defendant was arrested by the police one year after the crime, what would be your level of emotional distress?

Third Party

5. If you were the judge in this case and the defendant was arrested by the police on the day of the crime, what level of punishment do you believe should be imposed on the defendant?

6. If you were the judge in this case and the defendant was arrested by the police on the day of the crime, what would be your attitude towards the defendant's sentence?

7. If you were the judge in this case and the defendant was arrested by the police on the day of the crime, what would be your level of emotional distress?

8. If you were the judge in this case and the defendant was arrested by the police one year after the crime, what level of punishment do you believe should be imposed on the defendant?

9. If you were the judge in this case and the defendant was arrested by the police one year after the crime, what would be your attitude towards the defendant's sentence?

10. If you were the judge in this case and the defendant was arrested by the police one year after the crime, what would be your level of emotional distress?

(2) On May 20, 2017, the defendant, Yuan, decided to rob the person living next door. Yuan entered Hu's house through the window while brandishing a knife. Upon being startled awake and yelling, Hu was stabbed by Yuan. After Hu pleaded for mercy and offered to give Yuan money, Yuan stopped stabbing him, took the money, and fled the scene. Hu's injuries were classified as a Level one disability.

Second Party

1. If you were the victim in this case and the defendant was arrested by the police on the day of the crime, what level of punishment do you believe should be imposed on the defendant?

2. If you were the victim in this case and the defendant was arrested by the police on the day of the crime, what would be your level of emotional distress?

3. If you were the victim in this case and the defendant was arrested by the police three years later, what level of punishment do you believe should be imposed on the defendant?

4. If you were the victim in this case and the defendant was arrested by the police three years later, what would be your level of emotional distress?

Third Party

5. If you were the judge in this case and the defendant was arrested by the police on the day of the crime, what level of punishment do you believe should be imposed on the defendant?

6. If you were the judge in this case and the defendant was arrested by the police on the day of the crime, what would be your attitude towards the defendant's sentence?

7. If you were the judge in this case and the defendant was arrested by the police on the day of the crime, what would be your level of emotional distress?

8. If you were the judge in this case and the defendant was arrested by the police three years later, what level of punishment do you believe should be imposed on the defendant?

9. If you were the judge in this case and the defendant was arrested by the police three years later, what would be your attitude towards the defendant's sentence?

10. If you were the judge in this case and the defendant was arrested by the police three years later, what would be your level of emotional distress?

(3) On October 26, 2009, while residing in a village in Pu'an County, the defendant, Xie, was burning weeds on his property. Due to the dry weather conditions, the thatch surrounding the fire ignited and rapidly spread, causing a forest fire on the hill where the victim's family had rented land. Upon realizing that he could not extinguish the fire using branches, Xie immediately decided to return home. The Pu'an County Forest Fire Prevention Office reported the fire to the police on the same day. Upon investigation, it was determined that the fire caused economic losses of approximately 300,000 yuan to the victim, Li.

Second Party

1. If you were the victim in this case and the defendant was arrested by the police on the day of the crime, what level of punishment do you believe should be imposed on the defendant?

2. If you were the victim in this case and the defendant was arrested by the police on the day of the crime, what would be your level of emotional distress?

3. If you were the victim in this case and the defendant was arrested by the police nine years later, what level of punishment do you believe should be imposed on the defendant?

4. If you were the victim in this case and the defendant was arrested by the police nine years later, what would be your level of emotional distress?

Third Party

5. If you were the judge in this case and the defendant was arrested by the police on the day of the crime, what level of punishment do you believe should be imposed on the defendant?

6. If you were the judge in this case and the defendant was arrested by the police on the day of the crime, what would be your attitude towards the defendant's sentence?

7. If you were the judge in this case and the defendant was arrested by the police on the day of the crime, what would be your level of emotional distress?

8. If you were the judge in this case and the defendant was arrested by the police nine years later, what level of punishment do you believe should be imposed on the defendant?

9. If you were the judge in this case and the defendant was arrested by the police nine years later, what would be your attitude towards the defendant's sentence?

10. If you were the judge in this case and the defendant was arrested by the police nine years later, what would be your level of emotional distress?

(4) On May 8, 2018, the defendant, Kong, was informed by his wife that their son had accidentally hit a car parked in a fire lane while playing. Kong believed that the incident was due to the victim, Hu, incorrectly parking the car in the fire lane for an extended period. Subsequently, Kong used a key to damage the victim's car, causing extensive scratches. An appraisal determined that the car required 10,000 yuan in repairs. Following Hu's police report, the public security authorities initiated an investigation.

Second Party

1. If you were the victim in this case and the defendant was arrested by the police on the day of the crime, what level of punishment do you believe should be imposed on the defendant?

2. If you were the victim in this case and the defendant was arrested by the police on the day of the crime, what would be your level of emotional distress?

3. If you were the victim in this case and the defendant was arrested by the police two years later, what level of punishment do you believe should be imposed on the defendant?

4. If you were the victim in this case and the defendant was arrested by the police two years later, what would be your level of emotional distress?

Third Party

5. If you were the judge in this case and the defendant was arrested by the police on the day of the crime, what level of punishment do you believe should be imposed on the defendant?

6. If you were the judge in this case and the defendant was arrested by the police on the day of the crime, what would be your attitude towards the defendant's sentence?

7. If you were the judge in this case and the defendant was arrested by the police on the day of the crime, what would be your level of emotional distress?

8. If you were the judge in this case and the defendant was arrested by the police two years later, what level of punishment do you believe should be imposed on the defendant?

9. If you were the judge in this case and the defendant was arrested by the police two years later, what would be your attitude towards the defendant's sentence?

10. If you were the judge in this case and the defendant was arrested by the police two years later, what would be your level of emotional distress?

(5) On February 2, 2016, the defendant, Tan, threatened his ex-girlfriend, Liang, stating that he would post nude photographs of her on the internet to force her to meet him. Liang was compelled to meet Tan that evening, and Tan subsequently took her home on his motorcycle. Despite Liang's protests, Tan forced her to engage in sexual intercourse with him in his bed. The victim, Liang, then reported the incident to the police station. The police promptly dispatched officers to apprehend Tan; however, Tan fled, fearing further legal consequences.

Second Party

1. If you were the victim in this case and the defendant was arrested by the police on the day of the crime, what level of punishment do you believe should be imposed on the defendant?

2. If you were the victim in this case and the defendant was arrested by the police on the day of the crime, what would be your level of emotional distress?

3. If you were the victim in this case and the defendant was arrested by the police eight years later, what level of punishment do you believe should be imposed on the defendant?

4. If you were the victim in this case and the defendant was arrested by the police eight years later, what would be your level of emotional distress?

Third Party

5. If you were the judge in this case and the defendant was arrested by the police on the day of the crime, what level of punishment do you believe should be imposed on the defendant?

6. If you were the judge in this case and the defendant was arrested by the police on the day of the crime, what would be your attitude towards the defendant's sentence?

7. If you were the judge in this case and the defendant was arrested by the police on the day of the crime, what would be your level of emotional distress?

8. If you were the judge in this case and the defendant was arrested by the police eight years later, what level of punishment do you believe should be imposed on the defendant?

9. If you were the judge in this case and the defendant was arrested by the police eight years later, what would be your attitude towards the defendant's sentence?

10. If you were the judge in this case and the defendant was arrested by the police eight years later, what would be your level of emotional distress?

(6) On February 10, 2016, the defendant, Yuan, drove to the victim, Chen's home in a community in Wuhan City, under the assumption that residents near the river beach were affluent. Yuan used scissors to cut the door lock of Chen's house and stole numerous pieces of jewelry worth nearly one million dollars from Chen's safe. After committing the crime, Yuan fled the scene in a car with a false license plate. The victim, Chen, reported the theft to the public security authorities.

Second Party

1. If you were the victim in this case and the defendant was arrested by the police on the day of the crime, what level of punishment do you believe should be imposed on the defendant?

2. If you were the victim in this case and the defendant was arrested by the police on the day of the crime, what would be your level of emotional distress?

3. If you were the victim in this case and the defendant was arrested by the police four years later, what level of punishment do you believe should be imposed on the defendant?

4. If you were the victim in this case and the defendant was arrested by the police four years later, what would be your level of emotional distress?

Third Party

5. If you were the judge in this case and the defendant was arrested by the police on the day of the crime, what level of punishment do you believe should be imposed on the defendant?

6. If you were the judge in this case and the defendant was arrested by the police on the day of the crime, what would be your attitude towards the defendant's sentence?

7. If you were the judge in this case and the defendant was arrested by the police on the day of the crime, what would be your level of emotional distress?

8. If you were the judge in this case and the defendant was arrested by the police four years later, what level of punishment do you believe should be imposed on the defendant?

9. If you were the judge in this case and the defendant was arrested by the police four years later, what would be your attitude towards the defendant's sentence?

10. If you were the judge in this case and the defendant was arrested by the police four years later, what would be your level of emotional distress?

(7) On July 2, 2014, a minor dispute arose between the defendant, Xiao, and the victim, Zhao, while they were both employed at a factory in Guangzhou. During the altercation, the defendant struck Zhao's ribs with a bamboo pole, resulting in multiple fractures. Following a medical assessment, the victim was determined to have sustained a second-degree disability from his injuries. The defendant subsequently fled the scene and assumed a pseudonym.

Second Party

1. If you were the victim in this case and the defendant was arrested by the police on the day of the crime, what level of punishment do you believe should be imposed on the defendant?

2. If you were the victim in this case and the defendant was arrested by the police on the day of the crime, what would be your level of emotional distress?

3. If you were the victim in this case and the defendant was arrested by the police seven years later, what level of punishment do you believe should be imposed on the defendant?

4. If you were the victim in this case and the defendant was arrested by the police seven years later, what would be your level of emotional distress?

Third Party

5. If you were the judge in this case and the defendant was arrested by the police on the day of the crime, what level of punishment do you believe should be imposed on the defendant?

6. If you were the judge in this case and the defendant was arrested by the police on the day of the crime, what would be your attitude towards the defendant's sentence?

7. If you were the judge in this case and the defendant was arrested by the police on the day of the crime, what would be your level of emotional distress?

8. If you were the judge in this case and the defendant was arrested by the police seven years later, what level of punishment do you believe should be imposed on the defendant?

9. If you were the judge in this case and the defendant was arrested by the police seven years later, what would be your attitude towards the defendant's sentence?

10. If you were the judge in this case and the defendant was arrested by the police seven years later, what would be your level of emotional distress?

(8) On March 29, 2013, the defendant, Gao, took a taxi driven by the victim, Cheng. Noticing that Cheng was a woman, Gao planned a robbery. After the car arrived at an intersection in Puyang City, Gao threatened Cheng, demanding money, and stabbed her hands. He then held a fruit knife to her throat. Gao subsequently exited the vehicle and fled the scene. Cheng reported the incident to the police, and the public security agency initiated an investigation. Cheng's injuries resulted in a first-level disability.

Second Party

1. If you were the victim in this case and the defendant was arrested by the police on the day of the crime, what level of punishment do you believe should be imposed on the defendant?

2. If you were the victim in this case and the defendant was arrested by the police on the day of the crime, what would be your level of emotional distress?

3. If you were the victim in this case and the defendant was arrested by the police six years later, what level of punishment do you believe should be imposed on the defendant?

4. If you were the victim in this case and the defendant was arrested by the police six years later, what would be your level of emotional distress?

Third Party

5. If you were the judge in this case and the defendant was arrested by the police on the day of the crime, what level of punishment do you believe should be imposed on the defendant?

6. If you were the judge in this case and the defendant was arrested by the police on the day of the crime, what would be your attitude towards the defendant's sentence?

7. If you were the judge in this case and the defendant was arrested by the police on the day of the crime, what would be your level of emotional distress?

8. If you were the judge in this case and the defendant was arrested by the police six years later, what level of punishment do you believe should be imposed on the defendant?

9. If you were the judge in this case and the defendant was arrested by the police six years later, what would be your attitude towards the defendant's sentence?

10. If you were the judge in this case and the defendant was arrested by the police six years later, what would be your level of emotional distress?

(9) On July 29, 2010, the defendant, Niu, a kindergarten teacher with a car, drove to a village near Kaifeng City to pick up students. After loading the school bus with children, including the victim, Chen, Niu returned to the kindergarten. Upon arrival at the kindergarten, Niu disembarked and led the children to their respective classrooms. The victim, Chen, died from heatstroke due to the high temperature inside the school bus, as Niu neglected to check the bus after disembarking. Subsequently, Niu fled the scene out of fear of legal consequences.

Second Party

1. If you were the victim in this case and the defendant was arrested by the police on the day of the crime, what level of punishment do you believe should be imposed on the defendant?

2. If you were the victim in this case and the defendant was arrested by the police on the day of the crime, what would be your level of emotional distress?

3. If you were the victim in this case and the defendant was arrested by the police five years later, what level of punishment do you believe should be imposed on the defendant?

4. If you were the victim in this case and the defendant was arrested by the police five years later, what would be your level of emotional distress?

Third Party

5. If you were the judge in this case and the defendant was arrested by the police on the day of the crime, what level of punishment do you believe should be imposed on the defendant?

6. If you were the judge in this case and the defendant was arrested by the police on the day of the crime, what would be your attitude towards the defendant's sentence?

7. If you were the judge in this case and the defendant was arrested by the police on the day of the crime, what would be your level of emotional distress?

8. If you were the judge in this case and the defendant was arrested by the police five years later, what level of punishment do you believe should be imposed on the defendant?

9. If you were the judge in this case and the defendant was arrested by the police five years later, what would be your attitude towards the defendant's sentence?

10. If you were the judge in this case and the defendant was arrested by the police five years later, what would be your level of emotional distress?

Capital offense

(1) On March 30, 2007, the defendant Yu, facing financial difficulties, conceived the idea of kidnapping and extortion. Yu observed the victim Xiao (female, 13 years old at the time of death) walking alone and deceived her into entering a house under the pretense of helping with a task. Inside the house, Yu sexually assaulted Xiao. Subsequently, Yu departed the premises to demand a ransom from Xiao’s parents. Upon learning that Xiao’s family was actively searching for her, Yu became concerned about the potential exposure of his actions and decided to murder Xiao. He wrapped tape around her head and face and suffocated her using a quilt, leading to her death by mechanical asphyxiation.

Second Party

1. If you were the victim in this case and the defendant was arrested by the police on the day of the crime, what level of punishment do you believe should be imposed on the defendant?

2. If you were the victim in this case and the defendant was arrested by the police on the day of the crime, what would be your level of emotional distress?

3. If you were the victim in this case and the defendant was arrested by the police twelve years after the crime, what level of punishment do you believe should be imposed on the defendant?

4. If you were the victim in this case and the defendant was arrested by the police twelve years after the crime, what would be your level of emotional distress?

Third Party

5. If you were the victim in this case and the defendant was arrested by the police on the day of the crime, what level of punishment do you believe should be imposed on the defendant?

6. If you were the judge in this case and the defendant was arrested by the police on the day of the crime, what level of punishment do you believe should be imposed on the defendant?

7. If you were the judge in this case and the defendant was arrested by the police on the day of the crime, what would be your level of emotional distress?

8. If you were the judge in this case and the defendant was arrested by the police twelve years after the crime, what level of punishment do you believe should be imposed on the defendant?

9. If you were the judge in this case and the defendant was arrested by the police twelve years after the crime, what would be your stance on the defendant’s sentence?

10. If you were the judge in this case and the defendant was arrested by the police twelve years after the crime, what would be your level of emotional distress?

(2) On June 17, 2011, the defendant Ma encountered the victim Ren (female, aged 11) in the residential building where he lived as she was returning home from school. Under the pretext of helping his son with his homework, Ma lured her to his home, where he indecently assaulted her. Fearing that his actions would be exposed, Ma strangled Ren with a rope around her neck, causing her death by mechanical asphyxiation. Subsequently, the defendant Ma left his home and fled the scene.

Second Party

1. If you were the victim in this case and the defendant was arrested by the police on the day of the crime, what level of punishment do you believe should be imposed on the defendant?

2. If you were the victim in this case and the defendant was arrested by the police on the day of the crime, what would be your level of emotional distress?

3. If you were the victim in this case and the defendant was arrested by the police three years after the crime, what level of punishment do you believe should be imposed on the defendant?

4. If you were the victim in this case and the defendant was arrested by the police three years after the crime, what would be your level of emotional distress?

Third Party

5. If you were the judge in this case and the defendant was arrested by the police on the day of the crime, what level of punishment do you believe should be imposed on the defendant?

6. If you were the judge in this case and the defendant was arrested by the police on the day of the crime, what would be your attitude towards the defendant's sentence?

7. If you were the judge in this case and the defendant was arrested by the police on the day of the crime, what would be your level of emotional distress?

8. If you were the judge in this case and the defendant was arrested by the police three years after the crime, what level of punishment do you believe should be imposed on the defendant?

9. If you were the judge in this case and the defendant was arrested by the police three years after the crime, what would be your attitude towards the defendant's sentence?

10. If you were the judge in this case and the defendant was arrested by the police three years after the crime, what would be your level of emotional distress?

(3) On March 24, 2012, the defendant Luan, in an attempt to fraudulently obtain insurance money, placed a pre-purchased pesticide into the porridge consumed by his mother, Li (the victim), resulting in her death. Subsequently, Luan intended to use his status as Li's heir to fraudulently claim approximately 100,000 yuan in death insurance benefits from the insurance company. The public security organs determined Luan's guilt during the claims process but discovered that he had absconded.

Second Party

1. If you were the victim in this case and the defendant was arrested by the police on the day of the crime, what level of punishment do you believe should be imposed on the defendant?

2. If you were the victim in this case and the defendant was arrested by the police on the day of the crime, what would be your level of emotional distress?

3. If you were the victim in this case and the defendant was arrested by the police six years later, what level of punishment do you believe should be imposed on the defendant?

4. If you were the victim in this case and the defendant was arrested by the police six years later, what would be your level of emotional distress?

Third Party

5. If you were the judge in this case and the defendant was arrested by the police on the day of the crime, what level of punishment do you believe should be imposed on the defendant?

6. If you were the judge in this case and the defendant was arrested by the police on the day of the crime, what would be your attitude towards the defendant's sentence?

7. If you were the judge in this case and the defendant was arrested by the police on the day of the crime, what would be your level of emotional distress?

8. If you were the judge in this case and the defendant was arrested by the police six years later, what level of punishment do you believe should be imposed on the defendant?

9. If you were the judge in this case and the defendant was arrested by the police six years later, what would be your attitude towards the defendant's sentence?

10. If you were the judge in this case and the defendant was arrested by the police six years later, what would be your level of emotional distress?

(4) On January 21, 2003, the defendant Du planned to take revenge on the victim Deng due to Deng's continuous criticism of the civil engineering work performed by Du's company on the track. Du had prepared hypnotic medication in advance and added it to Deng's drink. After Deng fell asleep, Du struck Deng's head with a rubber mallet, causing severe brain injury and instant death. Du then transported Deng's body to a pre-selected field location and buried it in a pit that same night.

Second Party

1. If you were the victim in this case and the defendant was arrested by the police on the day of the crime, what level of punishment do you believe should be imposed on the defendant?

2. If you were the victim in this case and the defendant was arrested by the police on the day of the crime, what would be your level of emotional distress?

3. If you were the victim in this case and the defendant was arrested by the police fifteen years later, what level of punishment do you believe should be imposed on the defendant?

4. If you were the victim in this case and the defendant was arrested by the police fifteen years later, what would be your level of emotional distress?

Third Party

5. If you were the judge in this case and the defendant was arrested by the police on the day of the crime, what level of punishment do you believe should be imposed on the defendant?

6. If you were the judge in this case and the defendant was arrested by the police on the day of the crime, what would be your attitude towards the defendant's sentence?

7. If you were the judge in this case and the defendant was arrested by the police on the day of the crime, what would be your level of emotional distress?

8. If you were the judge in this case and the defendant was arrested by the police fifteen years later, what level of punishment do you believe should be imposed on the defendant?

9. If you were the judge in this case and the defendant was arrested by the police fifteen years later, what would be your attitude towards the defendant's sentence?

10. If you were the judge in this case and the defendant was arrested by the police fifteen years later, what would be your level of emotional distress?

(5) On October 15, 1999, the defendant Huang, facing financial difficulties, planned to rob the victim Xiao. Huang deceived Xiao by calling and pretending to introduce a business opportunity. He then drove Xiao's car to a remote road section in Fujian Province, where he used an electric shock baton to render Xiao unconscious. After robbing Xiao of cash and bank cards, Huang tied him up and subsequently strangled Xiao's neck with a belt, causing his death by mechanical asphyxiation. Huang then drove the car to the summit of a mountain and hid Xiao's body. Xiao's family reported him missing to the police.

Second Party

1. If you were the victim in this case and the defendant was arrested by the police on the day of the crime, what level of punishment do you believe should be imposed on the defendant?

2. If you were the victim in this case and the defendant was arrested by the police on the day of the crime, what would be your level of emotional distress?

3. If you were the victim in this case and the defendant was arrested by the police twenty-one years later, what level of punishment do you believe should be imposed on the defendant?

4. If you were the victim in this case and the defendant was arrested by the police twenty-one years later, what would be your level of emotional distress?

Third Party

5. If you were the judge in this case and the defendant was arrested by the police on the day of the crime, what level of punishment do you believe should be imposed on the defendant?

6. If you were the judge in this case and the defendant was arrested by the police on the day of the crime, what would be your attitude towards the defendant's sentence?

7. If you were the judge in this case and the defendant was arrested by the police on the day of the crime, what would be your level of emotional distress?

8. If you were the judge in this case and the defendant was arrested by the police twenty-one years later, what level of punishment do you believe should be imposed on the defendant?

9. If you were the judge in this case and the defendant was arrested by the police twenty-one years later, what would be your attitude towards the defendant's sentence?

10. If you were the judge in this case and the defendant was arrested by the police twenty-one years later, what would be your level of emotional distress?

(6) On October 27, 1995, the defendant Guan had a dispute with others on a farm in Suzhou City due to gambling. In an attempt to vent his rage, Guan drove an unlicensed car toward a group of people near the farm. The victim, Wang, was struck by Guan's car, causing his body to bounce off the ground and collide with the vehicle. As a result, Wang immediately lost consciousness. Guan immediately turned the car around and drove toward the crowd again, but the crowd managed to avoid him. Guan then fled the scene. The victim, Wang, died from a craniocerebral injury caused by the head impact.

Second Party

1. If you were the victim in this case and the defendant was arrested by the police on the day of the crime, what level of punishment do you believe should be imposed on the defendant?

2. If you were the victim in this case and the defendant was arrested by the police on the day of the crime, what would be your level of emotional distress?

3. If you were the victim in this case and the defendant was arrested by the police twenty-four years later, what level of punishment do you believe should be imposed on the defendant?

4. If you were the victim in this case and the defendant was arrested by the police twenty-four years later, what would be your level of emotional distress?

Third Party

5. If you were the judge in this case and the defendant was arrested by the police on the day of the crime, what level of punishment do you believe should be imposed on the defendant?

6. If you were the judge in this case and the defendant was arrested by the police on the day of the crime, what would be your attitude towards the defendant's sentence?

7. If you were the judge in this case and the defendant was arrested by the police on the day of the crime, what would be your level of emotional distress?

8. If you were the judge in this case and the defendant was arrested by the police twenty-four years later, what level of punishment do you believe should be imposed on the defendant?

9. If you were the judge in this case and the defendant was arrested by the police twenty-four years later, what would be your attitude towards the defendant's sentence?

10. If you were the judge in this case and the defendant was arrested by the police twenty-four years later, what would be your level of emotional distress?

(7) On October 6, 2009, a dispute erupted between the defendant Yang and his spouse, the victim Zhang, in the employee dormitory of Fuqing City. Zhang suspected that Yang was engaging in inappropriate relationships with other women at the company. In a fit of rage, Yang grabbed Zhang's neck with both hands in the bedroom, struggled with her, and then called for assistance for approximately two minutes, causing Zhang to die immediately. Zhang was identified as having been strangled, resulting in mechanical asphyxia and death. Yang then fled the crime scene.

Second Party

1. If you were the victim in this case and the defendant was arrested by the police on the day of the crime, what level of punishment do you believe should be imposed on the defendant?

2. If you were the victim in this case and the defendant was arrested by the police on the day of the crime, what would be your level of emotional distress?

3. If you were the victim in this case and the defendant was arrested by the police eighteen years later, what level of punishment do you believe should be imposed on the defendant?

4. If you were the victim in this case and the defendant was arrested by the police eighteen years later, what would be your level of emotional distress?

Third Party

5. If you were the judge in this case and the defendant was arrested by the police on the day of the crime, what level of punishment do you believe should be imposed on the defendant?

6. If you were the judge in this case and the defendant was arrested by the police on the day of the crime, what would be your attitude towards the defendant's sentence?

7. If you were the judge in this case and the defendant was arrested by the police on the day of the crime, what would be your level of emotional distress?

8. If you were the judge in this case and the defendant was arrested by the police eighteen years later, what level of punishment do you believe should be imposed on the defendant?

9. If you were the judge in this case and the defendant was arrested by the police eighteen years later, what would be your attitude towards the defendant's sentence?

10. If you were the judge in this case and the defendant was arrested by the police eighteen years later, what would be your level of emotional distress?

(8) On September 16, 2013, the victim, Wang, reprimanded the defendant Huang for poor internal management. Huang considered killing Wang, believing that Wang had deliberately retaliated against him and that the punishment was unjust. Later that afternoon, Huang entered Wang's office while Wang was seated unprepared in a chair. Huang then struck Wang's head multiple times with an axe before fleeing the scene. Wang's cause of death was determined to be a craniocerebral injury and a cervical fracture.

Second Party

1. If you were the victim in this case and the defendant was arrested by the police on the day of the crime, what level of punishment do you believe should be imposed on the defendant?

2. If you were the victim in this case and the defendant was arrested by the police on the day of the crime, what would be your level of emotional distress?

3. If you were the victim in this case and the defendant was arrested by the police nine years later, what level of punishment do you believe should be imposed on the defendant?

4. If you were the victim in this case and the defendant was arrested by the police nine years later, what would be your level of emotional distress?

Third Party

5. If you were the judge in this case and the defendant was arrested by the police on the day of the crime, what level of punishment do you believe should be imposed on the defendant?

6. If you were the judge in this case and the defendant was arrested by the police on the day of the crime, what would be your attitude towards the defendant's sentence?

7. If you were the judge in this case and the defendant was arrested by the police on the day of the crime, what would be your level of emotional distress?

8. If you were the judge in this case and the defendant was arrested by the police nine years later, what level of punishment do you believe should be imposed on the defendant?

9. If you were the judge in this case and the defendant was arrested by the police nine years later, what would be your attitude towards the defendant's sentence?

10. If you were the judge in this case and the defendant was arrested by the police nine years later, what would be your level of emotional distress?

(9) On October 15, 1990, the defendant, Cha, and the victim, Wang, were having a conversation over tea at a restaurant. Unbeknownst to the onlookers, the two men engaged in a physical altercation, pushing and shoving each other. After verbally threatening Wang, Cha returned to his house, retrieved his gun, and went in search of Wang. Upon encountering Wang in a building, Cha shot him multiple times in various parts of his body. Wang died a route to the hospital from the gunshot wounds. After committing the crime, Cha returned home with the shotgun and then fled. Wang was identified as having died from severe blood loss due to an organ rupture.

Second Party

1. If you were the victim in this case and the defendant was arrested by the police on the day of the crime, what level of punishment do you believe should be imposed on the defendant?

2. If you were the victim in this case and the defendant was arrested by the police on the day of the crime, what would be your level of emotional distress?

3. If you were the victim in this case and the defendant was arrested by the police twenty-seven years later, what level of punishment do you believe should be imposed on the defendant?

4. If you were the victim in this case and the defendant was arrested by the police twenty-seven years later, what would be your level of emotional distress?

Third Party

5. If you were the judge in this case and the defendant was arrested by the police on the day of the crime, what level of punishment do you believe should be imposed on the defendant?

6. If you were the judge in this case and the defendant was arrested by the police on the day of the crime, what would be your attitude towards the defendant's sentence?

7. If you were the judge in this case and the defendant was arrested by the police on the day of the crime, what would be your level of emotional distress?

8. If you were the judge in this case and the defendant was arrested by the police twenty-seven years later, what level of punishment do you believe should be imposed on the defendant?

9. If you were the judge in this case and the defendant was arrested by the police twenty-seven years later, what would be your attitude towards the defendant's sentence?

10. If you were the judge in this case and the defendant was arrested by the police twenty-seven years later, what would be your level of emotional distress?
